# The histone demethylase KDM5 has insulator activity in the brain

**DOI:** 10.1101/2024.12.04.626780

**Authors:** Matanel Yheskel, Melissa A. Castiglione, Richard D. Kelly, Simone Sidoli, Julie Secombe

## Abstract

KDM5 family proteins are best known for their demethylation of the promoter proximal chromatin mark H3K4me3. KDM5-regulated transcription is critical in the brain, with variants in the X-linked paralog *KDM5C* causing the intellectual disability (ID) disorder Claes-Jensen syndrome. Although the demethylase activity of KDM5C is known to be important for neuronal function, the contribution of non-enzymatic activities remain less characterized. We therefore used *Drosophila* to model the ID variant *Kdm5^L854F^*, which disrupts a C5HC2 zinc finger adjacent to the enzymatic JmjC domain. *Kdm5^L854F^* causes similar transcriptional changes in the brain to a demethylase dead strain, *Kdm5^J1310C^\**, despite having little effect on enzymatic activity. KDM5^L854F^ is also distinct from KDM5^J1310C^* in its reduced interactions with insulator proteins and enhancement of position effect variegation. Instead, the common transcriptional deficits likely result from both the JmjC and C5HC2 domains driving proper genomic organization through their activity in promoting proper loop architecture.

## Introduction

The KDM5 family of enzymes are multidomain proteins that function as transcriptional regulators. Across species, KDM5 proteins are expressed broadly across all cell types examined, but their function appears to be particularly important in neurons for brain development and function. Emphasizing this, genetic disruption to three of the four human *KDM5* paralogs, *KDM5A*, *KDM5B*, and *KDM5C*, are found in individuals with neurodevelopmental disorders characterized by intellectual disability (ID), seizures and changes to behavior and movement.[1–4] The important role that KDM5 proteins play in neurons is evolutionarily conserved, with genetic knockout of *Kdm5a*, *Kdm5b,* or *Kdm5c* in *Mus musculus* causing altered learning and memory, social interactions, and seizure susceptibility.[5–8] Invertebrates such as *Drosophila melanogaster* that encode a single KDM5 protein also show neuronal phenotypes that are consistent with those observed in humans. These include KDM5-dependent functions in learning and memory, neurotransmission, seizure-like activity and locomotion.[9–11]

*KDM5C* is the best described paralog with respect to its link to neurodevelopmental disorders, with pathogenic alleles that cause Claes-Jensen X-linked Intellectual disability syndrome representing a wide range of loss of function nonsense, missense, frame shift, and splice site changes.[1,7,12,13] Though nonsense variants in *KDM5C* result in lack of protein in hemizygous males, many missense variants do not appreciably affect protein stability.[1,13] These single amino acid changes occur across the KDM5C protein, including in the Jumonji C (JmjC) domain that functions with JmjN to form the active enzymatic site of KDM5 proteins.[1,3] The combined activity of the JmjN/C domains demethylates trimethylated lysine 4 of histone H3 (H3K4me3), a promoter-proximal histone modification associated with transcriptional activation and consistency.[14–16] In keeping with a key neuronal role for this demethylase activity, some cognitive and behavioral phenotypes of *Kdm5c* knockout mice can be partially suppressed by genetically restoring levels of H3K4me3.[17] In addition, analyses of a JmjC domain mutant *Drosophila* strain lacking demethylase activity (*Kdm5^JmjC*^*) displayed altered learning and memory, seizure susceptibility, locomotion and neurotransmission.[9,11] Despite this, the role that loss of enzymatic activity plays in the pathogenesis of Claes-Jensen syndrome remains unclear. Indeed, many missense variants occur outside of the enzymatic JmjN/JmjC domains, and several of these have been shown to have only slight or no effect on demethylase activity *in vitro*.[1,13] These data suggest that demethylase-dependent and demethylase-independent effects on gene expression contribute to the phenotypes observed in those with Claes-Jensen syndrome. This is consistent with a broader body of literature regarding KDM5 proteins showing that they possess demethylase-independent gene regulatory functions.[1,9,10,18–24] These include effects on histone acetylation, promoter-proximal pausing, and nucleosome positioning.[15,22] When and where these non-canonical KDM5 activities are utilized *in vivo*, their relationship to the histone demethylase activity, and whether KDM5 proteins can also impact gene regulation through additional activities, remains unknown. ID-associated alleles therefore provide us with an opportunity to uncover new gene-regulatory activities of KDM5 proteins, and to provide insights into the links between KDM5 and neuronal (dys)function.

Here we use *Drosophila* to investigate the molecular deficits of *KDM5C^L731F^*, an ID allele associated with severe intellectual disability, ataxia and increased aggression.[25,26] This missense variant occurs within the C5HC2 zinc finger that has no known function, and results in a 2-fold decrease to *in vitro* enzymatic activity.[27] However, the molecular deficits of this variant protein in an *in vivo* context have not been investigated in detail.[27] We generated flies harboring an allele equivalent to *KDM5C^L731F^*in *Kdm5* (*Kdm5^L854F^*) as part of a library of human ID-allele variant strains.[9–11] Consistent with phenotypes observed in humans, *Kdm5^L854F^* adult flies show deficits in associative memory and altered locomotion.[9,10] However, in contrast to *Kdm5^JmjC*^* that lacks demethylase activity, *Kdm5^L854F^* does not significantly alter bulk levels of H3K4me3, suggesting that the KDM5^L854F^ variant protein retains some, or all, of its enzymatic activity *in vivo*. Consistent with this observation, this allele is phenotypically distinct from *Kdm5^JmjC*^* at the larval neuromuscular junction (NMJ), with *Kdm5^JmjC*^* but not *Kdm5^L854F^*, altering electrophysiological properties at this synapse.[10] Here we show that the changes to gene expression observed in the adult brain of *Kdm5^L854F^* animals are not explained by changes to H3K4me3. Instead, proximity-mediated studies reveal that KDM5^L854F^ displays reduced proximity to several previously identified KDM5 interactors, particularly with known insulators. Consistent with playing key roles in genomic organization, KDM5 is found at loop anchors, and this correlates with altered gene expression in *Kdm5^L854F^* and in *Kdm5^JmjC*^*. This suggests that the JmjC and C5HC2 domains are both necessary at these sites, and that this may explain the similar changes to gene expression seen in *Kdm5^L854F^*and *Kdm5^JmjC*^*. Combined, our analyses of an ID-associated allele have revealed new functions for KDM5 in genomic structure and has provided new insights into the pathogenic mechanisms that may contribute to Claes-Jensen syndrome.

## Results

### *Kdm5^L854F^* shows similar changes to gene expression to *Kdm5^JmjC*^*

The ID-associated *KDM5C^L731F^* variant occurs within a C5HC2 zinc finger domain that, consistent with its importance, is conserved in human KDM5A, KDM5B, KDM5C and KDM5D in addition to *Drosophila* KDM5 (Fig. 1A, B). The *Kdm5^L854F^* allele produces wild-type levels of protein and does not significantly alter bulk levels of H3K4me3 in the adult head, suggesting that it does not abolish enzymatic activity (Fig. 1C). To understand the contributions of demethylase-dependent and independent activities to KDM5-regulated transcription we therefore focused on comparing *Kdm5^L854F^* to a demethylase-dead strain (*Kdm5^JmjC*^*).[9–11] To examine the spatial relationship of the residues altered in the KDM5^JmjC*^ and KDM5^L854^ proteins, we modeled *Drosophila* KDM5 in AlphaFold (Fig. 1D).[28] Despite the introduction of a bulky phenylalanine residue into the C5HC2 domain in KDM5^L854^, there do not appear to be gross changes to the structure of this domain (Fig. 1E). Furthermore, Leucine 854 does not appear to interact with structural components necessary to maintain the demethylation binding pocket, as evidenced by the lack of a predicted change to the JmjC domain in KDM5^L854^, consistent with the possibility that KDM5^L854F^ retains enzymatic function (Fig. 1F).

**Figure 1:**
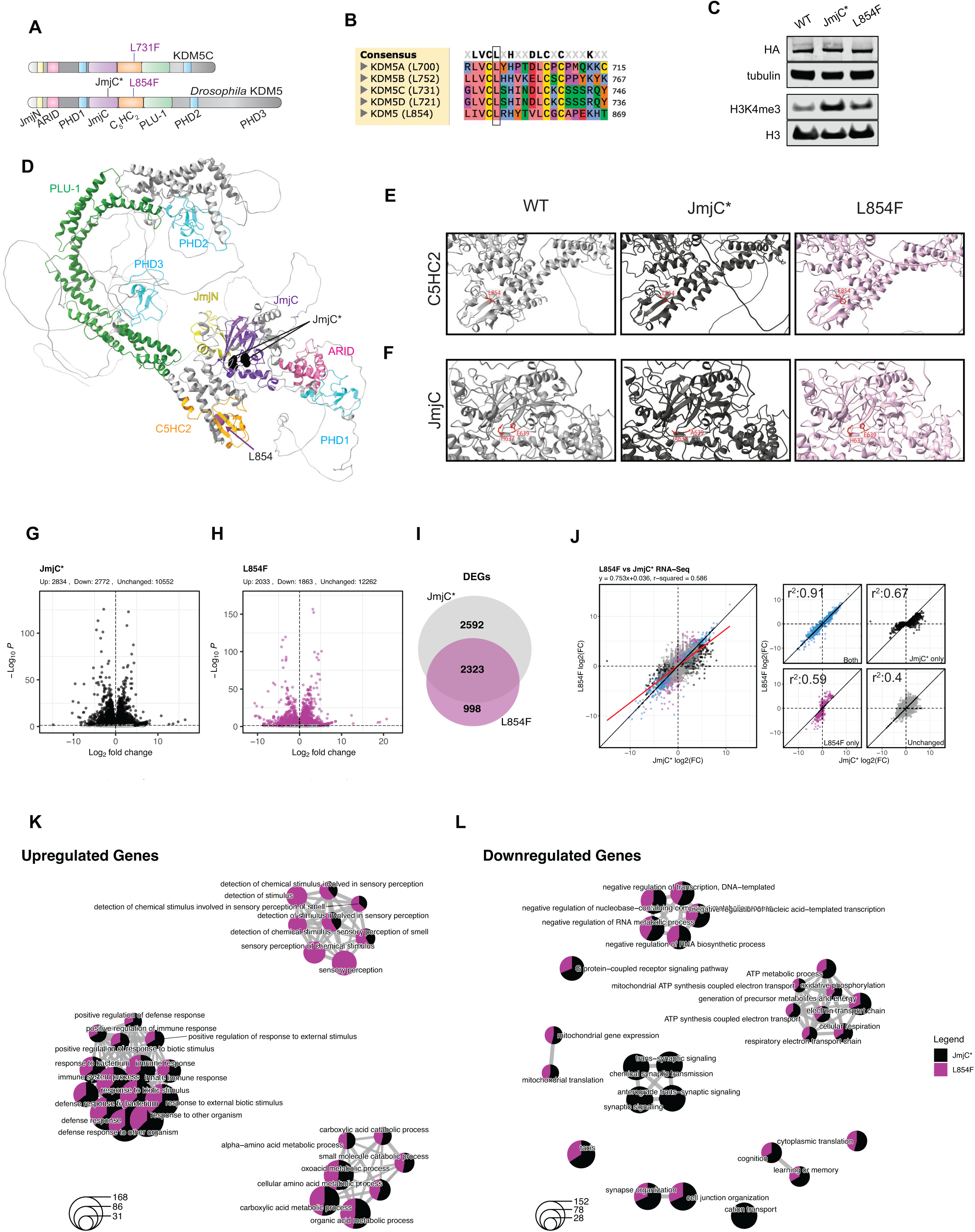
*Kdm5^JmjC*^* and *Kdm5^L854F^* show different effects on bulk H3K4me3 but similar overall changes to gene expression. A) Schematic of the ID-associated KDM5C^L731F^ variant and the analogous residue in *Drosophila* KDM5^L854F^. *Kdm5^JmjC*^* serves as a demethylase-dead control. B) Alignment of human KDM5A, KDM5B, KDM5C, KDM5D and *Drosophila* KDM5 showing amino acid conservation. Leucine 731 of KDM5C is conserved across all KDM5 proteins shown. C) Western blot analysis using adult heads from *Kdm5^WT^* (control), *Kdm5^JmjC*^*and *Kdm5^L854F^*. Shown are KDM5 detected using the HA tag, anti-tubulin loading control for anti-HA, anti-H3K4me3 and total H3. D) AlphaFold model of *Drosophila* KDM5 visualized using Chimera in which domains are indicated using the same color as in panel A. Bubbles represent the residues that are considered in this study. In black are the two residues which bind to the iron molecule necessary for demethylation (H637A and E639A; altered in KDM5^JmjC*^) and in purple is the L854F within the C5HC2 zinc finger. E) Closer view of C5HC2 domain in KDM5^WT^, KDM5^JmjC*^, and KDM5^L854F^ proteins. The side chain at 854 is shown and highlighted in red. F) Closer view of JmjC domain in KDM5^WT^, KDM5^JmjC*^, and KDM5^L854F^ proteins. The side chains altered in the demethylase dead KDM5^JmjC*^ protein, 637 and 639, are shown and highlighted in red. G) Volcano plot of RNA-seq data from *Kdm5^JmjC*^* adult heads. Black dots indicate significantly affected using an FDR cutoff of 5%. Grey indicates unchanged. H) Volcano plot of RNA-seq data from *Kdm5^L854F^* heads. Purple dots indicate significantly affected using an FDR cutoff of 5%. Grey indicates unchanged. I) Venn diagram showing how many differentially expressed genes (DEGs) overlap between *Kdm5^JmjC*^*and *Kdm5^L854F^*. J) Scatterplots of RNA-Seq log2(FC)s from *Kdm5^L854F^* and *Kdm5^JmjC*^*. Blue = DEG in both genotypes, black = DEG in *Kdm5^JmjC*^* only, purple = DEG in *Kdm5^L854F^* only and gray = unchanged in both. K) GO analysis of upregulated genes using clusterProfiler in which the size of each circle corresponds to the number of genes in the GO category and the size of each pie slice corresponds to the relative number of genes in each category attributed to that genotype according to color. L) GO analysis of downregulated genes using clusterProfiler.

We previously showed that *Kdm5^JmjC*^* and all ID-associated alleles examined to-date displayed a characteristic downregulation of genes required for the regulation of translation such as ribosomal protein genes in the adult brain.[9] Looking in greater depth at *Kdm5^L854F^* and *Kdm5^JmjC*^*, we compared all gene expression changes observed in these two genotypes. *Kdm5^JmjC*^*and *Kdm5^L854F^* showed similar numbers of up– and downregulated genes, with many of the differentially expressed genes (DEGs) overlapping (FDR<0.05; Fig. 1G, H; TableS1). Emphasizing their similarities, the expression of genes that were significantly dysregulated in both *Kdm5^JmjC*^* and *Kdm5^L854F^* animals were highly correlated (r^2^ = 0.91; p=2.2e-16; Fig. 1I, J). Consistent with this, *Kdm5^L854F^* and *Kdm5^JmjC*^* shared many gene-ontology biological process categories (GO-BP), although some were unique to one genotype or the other (Fig. 1K, L). In addition to the expected enrichment for ribosomal protein genes, other categories were shared between *Kdm5^L854F^* and *Kdm5^JmjC*^*.[9] These included categories related to cognition and synaptic organization, mitochondrial and other metabolic processes, and the regulation of transcription. Interestingly, *Kdm5^JmjC^\** additionally showed the downregulation of synaptic signaling and cation transport genes. These data suggest a role for the demethylase activity of KDM5 in maintaining the expression of these genes and is in keeping with electrophysiological phenotypes observed at the larval NMJ for *Kdm5^JmjC*^* and not *Kdm5^L854F^*.[10]

### *Kdm5^L854F^* and *Kdm5^JmjC*^* show distinct changes to the genomic distribution of H3K4me3

One possible explanation for the similarities in the gene expression changes observed in *Kdm5^L854F^*and *Kdm5^JmjC*^* animals is that despite the lack of change to overall levels of H3K4me3 in *Kdm5^L854F^*, some promoters exhibit a localized increase in this histone mark. To test this, we quantified genomic changes to H3K4me3 using <u>C</u>leavage <u>U</u>nder <u>T</u>argets & <u>R</u>elease <u>U</u>sing <u>N</u>uclease (CUT&RUN) using adult brains from *Kdm5^WT^*, *Kdm5^JmjC*^*, and *Kdm5^L854F^* animals. In *Kdm5^WT^* brains, 87% of H3K4me3 peaks were promoter proximal, and GO analyses of the associated genes revealed enrichment for terms associated with neuronal function (SFig. 1A-C). Consistent with the elevated overall level of H3K4me3 observed in *Kdm5^JmjC*^*animals by western blot, transcription start site (TSS) metagene profiles showed increased H3K4me3 in these animals compared to *Kdm5^WT^* (Fig 2A). In contrast, the overall profile of H3K4me3 in *Kdm5^L854F^* was similar to *Kdm5^WT^*, although a small increase can be seen downstream of the TSS suggesting that this allele may cause some minor changes to the distribution of this chromatin mark (Fig. 2A).

**Figure 2:**
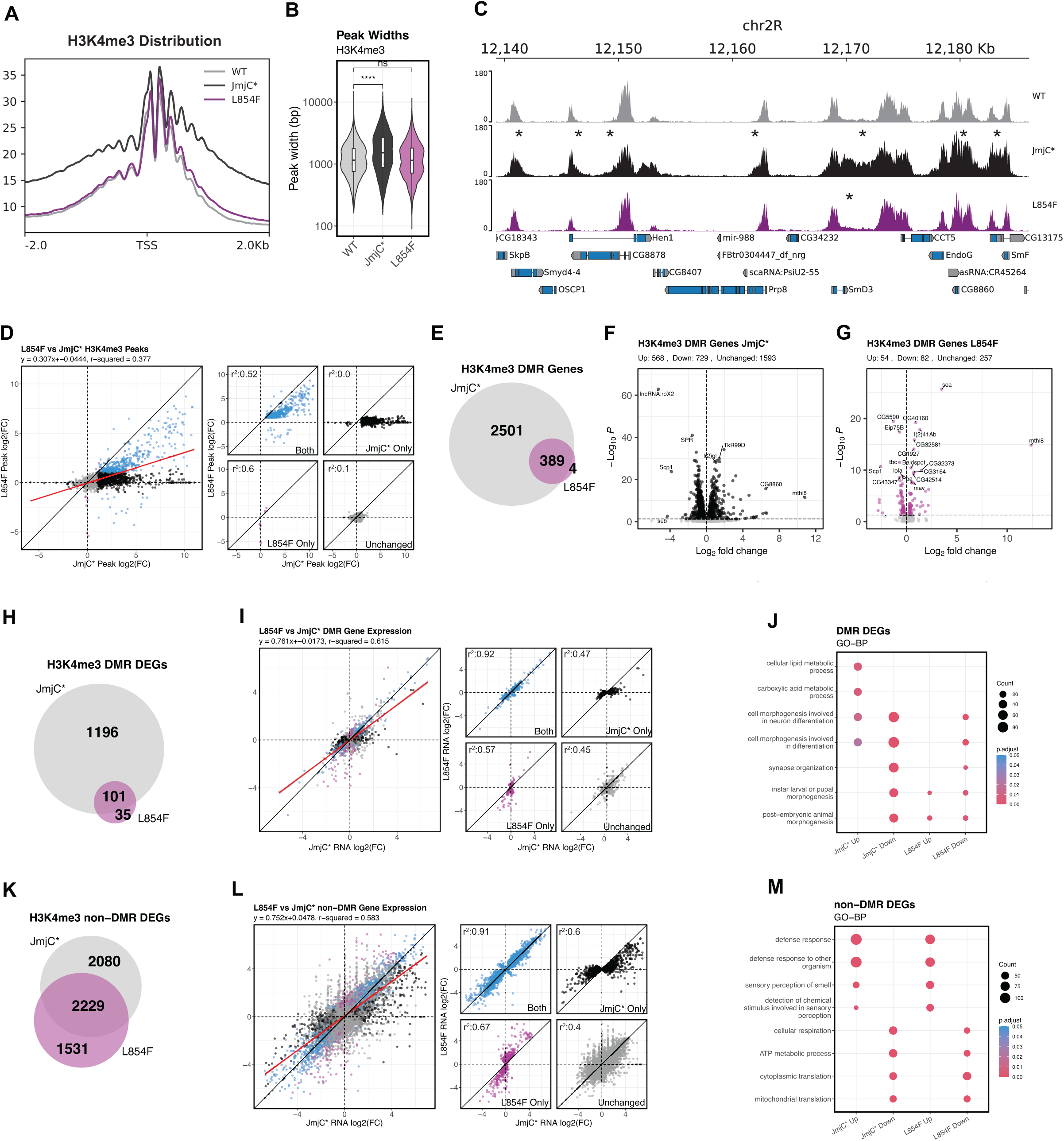
Increased promoter H3K4me3 does not correlate with changes to transcript levels. A) TSS profiles of H3K4me3 CUT&RUN from *Kdm5^WT^* (grey), *Kdm5^JmjC*^* (black), *Kdm5^L854F^* (purple). B) Quantitation of H3K4me3 peak widths from *Kdm5^WT^* (grey; median of 1152bp), *Kdm5^JmjC*^*(black; median of 1524bp), *Kdm5^L854F^* (purple; median of 1145). **** p-value<0.0001 Kruskal-Wallis test. C) Genome track of H3K4me3 CUT & RUN data from *Kdm5^WT^* (grey), *Kdm5^JmjC*^* (black), *Kdm5^L854F^* (purple). * indicates regions that show visually expanded regions of H3K4me3. D) Scatterplots of log2(FC) of H3K4me3 signal comparing *Kdm5^JmjC*^* to *Kdm5^WT^*. Blue = differentially methylated regions (DMRs) in both genotypes, purple = DMR in *Kdm5^L854F^* only, black = DMR in *Kdm5^JmjC*^* only, grey = peaks unchanged. Separated data shown on right. E) Venn diagram showing overlap between DMR-associated genes in both genotypes. F) Volcano plot showing changes to gene expression of DMR-associated genes in *Kdm5^JmjC*^* compared to *Kdm5^WT^*. G) Volcano plot showing changes to gene expression of DMR-associated genes in *Kdm5^L854F^* compared to *Kdm5^WT^*. H) Venn diagram showing overlap between DMR-associated DEGs for *Kdm5^JmjC*^* and *Kdm5^L854F^*. I) Scatterplot showing mRNA log2(FC)s of DMR-associated genes for *Kdm5^JmjC*^* and *Kdm5^L854F^*. Blue = DEGs in both genotypes, purple = DEGs in *Kdm5^L854F^* only, black = DEG in *Kdm5^JmjC*^* only, grey = peak unchanged. Separated data shown on right. J) GO analyses of DMR-associated DEGs in *Kdm5^L854F^* and *Kdm5^JmjC*^* separated by up– and downregulated genes. K) Venn diagram showing overlap between non DMR-associated DEGs in *Kdm5^JmjC*^* and *Kdm5^L854F^*. L) Scatterplot showing mRNA log2(FC)s of non DMR-associated genes in *Kdm5^JmjC*^* and *Kdm5^L854F^*. Blue = DEGs in both, purple = DEGs in *Kdm5^L854F^* only, black = DEGs in *Kdm5^JmjC*^* only, grey = peak unchanged. Separated data shown on right. M) GO analyses of non DMR-associated DEGs separated by genotype and by up– and downregulated genes.

The most striking change to H3K4me3 observed in *Kdm5^JmjC*^*animals was not to the height of the peak surrounding the TSS, but rather a spreading of this chromatin mark into gene bodies (Fig. 2A). Consistent with this, peak calling analyses using Genrich revealed that *Kdm5^JmjC*^* brains showed significantly wider H3K4me3 peaks compared to *Kdm5^L854F^* and *Kdm5^WT^* (p=2e-16; Fig. 2B, C). To define which changes to H3K4me3 were significantly altered in *Kdm5^JmjC*^* compared to *Kdm5^WT^*, we used DiffBind to identify differentially methylated regions (log2FC>1, FDR<0.01). *Kdm5^JmjC*^* brains showed more H3K4me3 at 2461 regions (40% of all peaks), consistent with KDM5 functioning broadly across the genome to regulate this chromatin mark (Fig. 2C indicated by *; Fig. S1D). In *Kdm5^L854F^* animals, while the average length of H3K4me3 peaks across the genome was not different from *Kdm5^WT^*, 433 regions had significantly increased H3K4me3, all but 10 of which were also identified in *Kdm5^JmjC*^* (Fig. 2C; SFig 1E-F). Although statistically significant, the changes to H3K4me3 in *Kdm5^L854F^* were much smaller than those seen in *Kdm5^JmjC*^* brains (Fig. 2C, D). The transcriptional similarities between the two alleles are therefore not caused by the variant KDM5^L854F^ protein lacking demethylase activity at a subset of promoters across the genome.

Peak annotation using ChIPSeeker identified 2890 genes with altered levels of H3K4me3 in *Kdm5^JmjC*^*, 389 of which were also observed in *Kdm5^L854F^* (Fig. 2E; TableS1).[29] These changes occurred primarily at the promoters of genes involved in neuron projection and morphogenesis, in addition to those encoding proteins localized to axons, vesicles, and synapses (SFig. 1G-I). However, changes to H3K4me3 levels did not correlate with altered mRNA levels detected by RNA-seq. More than half of genes with altered H3K4me3 in *Kdm5^JmjC*^*and *Kdm5^L854F^* animals showed no change to transcript levels, while the remaining genes were relatively evenly split between genes that were up-or downregulated (Fig. 2F, G; SFig. 1J-K). Only 101 genes showed increased H3K4me3 and similarly up-or downregulated transcript levels in *Kdm5^JmjC*^* and *Kdm5^L854F^*, including genes required for metabolic and neuronal functions (r^2^=0.92, blue dots; Fig. 2H-J). Interestingly, the group of genes that exhibited the strongest correlation were those that were differentially expressed in *Kdm5^JmjC*^* and *Kdm5^L854F^* but showed no significant change to promoter H3K4me3 (2229 genes; r^2^ = 0.91; p-value<2.2e-16; Fig. 2K, L). Genes in this category included those related to cytoplasmic translation such as ribosome protein genes, which are the most significant category of gene altered in *Kdm5^JmjC*^* and other ID-variant alleles (Fig. 2M).[9,11] Overall, these data show that the transcriptomic changes seen in *Kdm5^JmjC*^* and *Kdm5^L854F^* are not linked to changes to levels of promoter H3K4me3.

### Changes to H3K4me3 or gene expression are not driven by altered promoter recruitment of KDM5^JmjC*^ or KDM5^L854F^

To test whether changes to gene expression were due to altered KDM5 recruitment, we performed <u>Ta</u>rgeted <u>Da</u>mID (TaDa) in adult brains, comparing wildtype KDM5 to KDM5^JmjC*^ and KDM5^L854F^. To accurately reflect the proper cellular context, we pulsed on the expression of each Dam-KDM5 fusion protein or Dam alone control in their respective wildtype or *Kdm5* allele backgrounds during early adulthood. A total of 5,893 KDM5 binding sites were observed across all three genotypes, most of which were promoter proximal at genes related to neuronal development and function (Fig. 3A; SFig. 2A-C; TableS1). These TaDa data show KDM5 binding further downstream relative to the TSS compared to prior ChIP-seq data using whole adult flies. This could be due to differences in detection methods, as TaDa does not rely on stable KDM5 recruitment to genomic regions and does not utilize crosslinking.[30] Notably, these TaDa-derived genomic recruitment data coincide well with genomic regions that show extended H3K4me3 in *Kdm5^JmjC*^*, consistent with this reflecting a physiologically relevant location of KDM5 (Fig. 3B).[30–32] Comparing TaDa data from *Kdm5^JmjC*^* and *Kdm5^L854F^* to *Kdm5^WT^* revealed an overall decrease in peak height but extension of peak width surrounding the TSS (Fig. 3A-C). KDM5^JmjC*^ and KDM5^L854F^ can therefore be recruited to their target genes but show a slightly expanded distribution.

**Figure 3:**
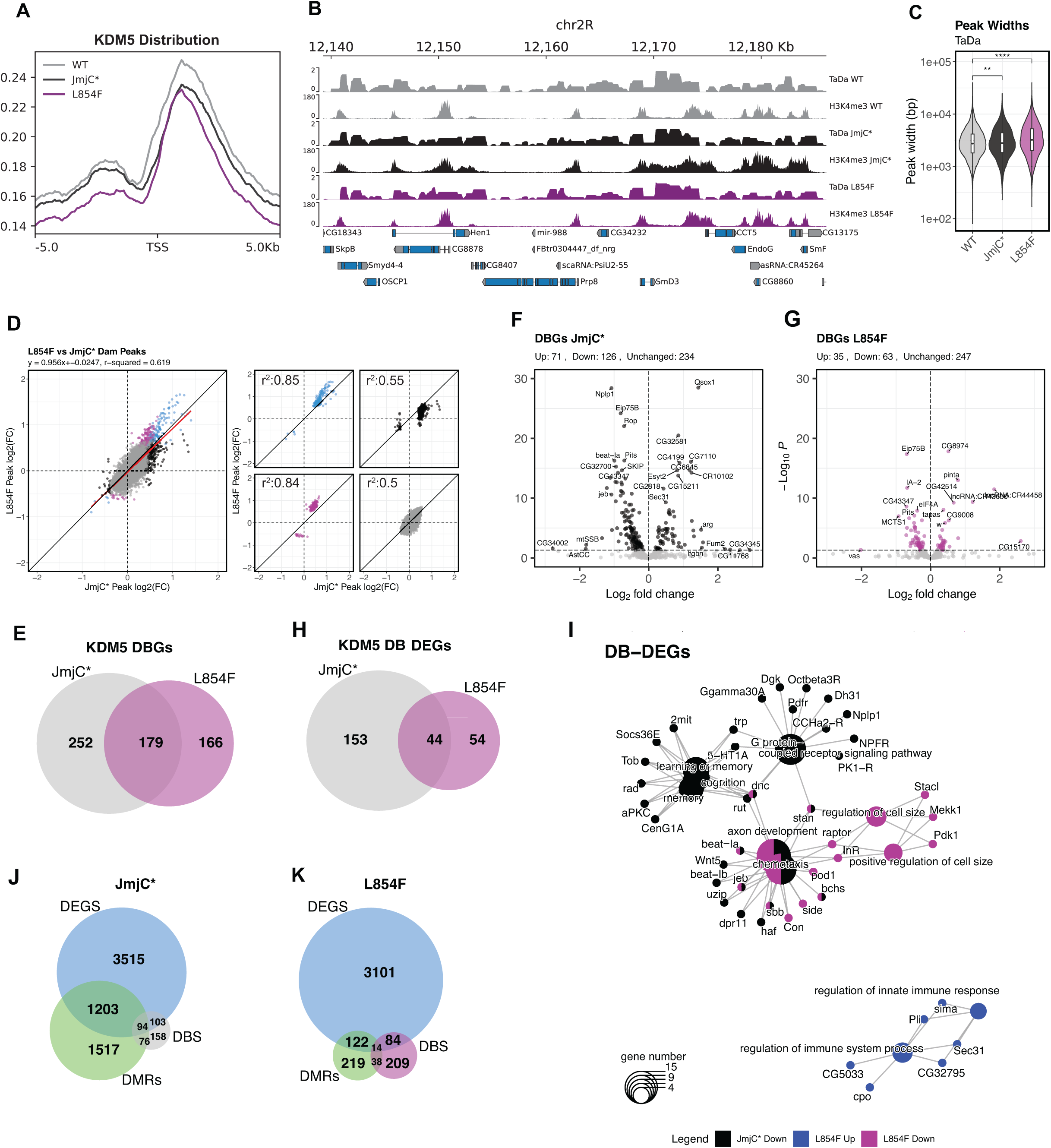
Targeted DamID (TaDa) shows occupancy of KDM5^WT^, KDM5^JmjC*^ and *Kdm5^L854F^*. A) TSS profiles of Dam:KDM5^WT^, Dam:KDM5^JmjC*^ and Dam:KDM5^L854F^ in adult heads after being pulsed on for 24 hours during early adulthood. B) Genome track of TaDa data from KDM5^WT^, KDM5^JmjC*^ and KDM5^L854F^ and matching H3K4me3 CUT&RUN data. C) TaDa peak widths. Median width KDM5^WT^ was 2727bp, 2788bp for KDM5^L854F^ and 3216bp for KDM5^JmjC*^. ** p-value < 0.01, **** p-value < 0.0001 D) Scatterplot of log2(FC) of H3K4me3 signal compared to wildtype. Blue = differentially bound sites (DBS) in both, purple = DBS in KDM5^L854F^ only, black = DBS in KDM5^JmjC*^ only, grey = peak unchanged. Separated data shown on right. E) Venn Diagram of differentially bound genes (DGBs) in both genotypes. F) Volcano plot showing gene expression changes of DGBs in *Kdm5^JmjC*^* compared to *Kdm5^WT^*. G) Volcano plot showing gene expression changes of DGBs in *Kdm5^L854F^* compared to *Kdm5^WT^*. H) Venn diagram showing genes associated with differentially bound regions in (DB DEGs) in *Kdm5^JmjC*^*and *Kdm5^L854F^*. I) GO analyses of genes that show altered binding and altered gene expression (DB DEGs). Black = decreased in *Kdm5^JmjC*^*, purple = decreased in *Kdm5^L854F^*, and blue = increased in *Kdm5^L854F^*. Lines denote the gene that contributes to the GO category. J) Venn Diagram comparing overlap of DMR-associated genes, DBGs, and DMRs in *Kdm5^JmjC*^*. K) Venn Diagram comparing overlap of DMR-associated genes, DBGs, and DMRs in *Kdm5^L854F^*.

To define the statistically significant changes to the KDM5 TaDa signal across genotypes we used NOISeq (log2FC>0.58 and p-value<0.2 cutoff).[33,34] KDM5^JmjC*^ and KDM5^L854F^ both displayed a slight increase in binding frequency or residence time at a subset of promoters across the genome (SFig. 2D-G). To link this to changes in mRNA levels detected by RNA-seq, we identified genes associated with altered KDM5 binding and found 431 differentially bound genes for *Kdm5^JmjC*^* and 345 for *Kdm5^L854F^*, 179 of which overlapped (Fig. 3E; TableS1). These genes were enriched for GO categories such as memory, cognition, and neuronal projection (SFig. 2H-I). Integrating the altered KDM5 binding and RNA-seq data did not identify a consistent theme. In both genotypes, while some genes that showed altered TaDa binding were either up-or downregulated, most were unchanged (54% and 72% for *Kdm5^JmjC*^* and *Kdm5^L854F^*, respectively; Fig. 3F, G). Of those genes that were transcriptionally dysregulated, 44 were shared between *Kdm5^JmjC*^* and *Kdm5^L854F^* consistent with these mutant proteins sharing some characteristics (Fig. 3H, I). Incorporating H3K4me3 CUT&RUN data with KDM5 binding and gene expression data revealed 94 genes in *Kdm5^JmjC*^* and 14 in *Kdm5^L854F^* with changes to all three categories (Fig. 3J, K; TableS1). Combined, these results suggest that variant KDM5 proteins have modestly increased binding to gene bodies/promoters. However, these changes are not likely to be driving the observed changes to H3K4me3 or to gene expression.

### KDM5^L854F^ and not KDM5^JmjC*^ shows reduced proximity to boundary element proteins

To determine if changes to gene expression seen in ID-variants could be explained through altered protein-protein interactions, we used *in vivo* TurboID-mediated proximity labeling. We previously used this method to describe a network of 87 high-confidence KDM5 interactors in the adult brain.[35] To examine the extent to which ID-variants affected this KDM5 interactome, we generated transgenic flies able to conditionally express N-terminal TurboID fusions of KDM5^JmjC*^ or KDM5^L854F^ (UASp-*TurboID*:*Kdm5^JmjC^\** and UASp-*TurboID*:*Kdm5^L854F^*). To enhance our ability to detect interactors, we created fly strains in which the sole source of KDM5 was TurboID:KDM5^variant^ by ubiquitously expressing these fusion proteins at approximately endogenous levels in a *Kdm5^Δ^* background using Ubi-Gal4, similar to our prior study of wild-type KDM5 (Fig. 4A).[35] As expected, expression of Turbo-tagged versions of *Kdm5^JmjC*^*or *Kdm5^L854F^* variant genes rescued the lethality of *Kdm5^Δ^*(SFig. 3A).[9]

**Figure 4:**
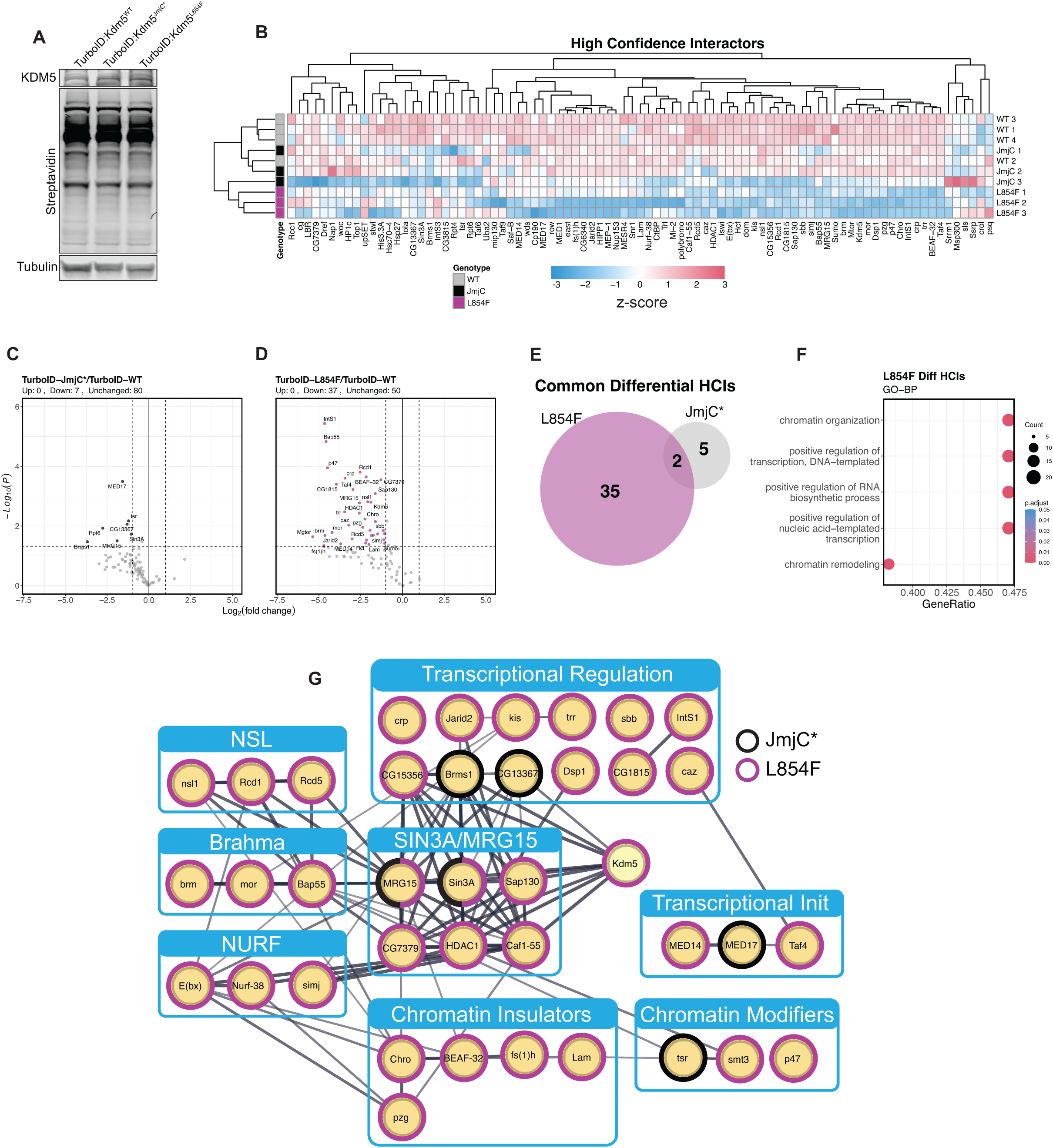
KDM5^L854F^ but not KDM5^JmjC*^ shows reduced proximity to insulator proteins. A) Western blots from adult heads expressing TurboID:KDM5^WT^, TurboID:KDM5^JmjC*^ or TurboID:KDM5^L854F^ heads. TurboID transgenes are UASp constructs and are expressed ubiquitously in a Kdm5^Δ^ background (*Kdm5^Δ^*, Ubi-Gal4/*Kdm5^Δ^*; UASp-TurboID:*Kdm5^WT^ ^JmjC*^ ^or^ ^L854F^*/+). Anti-KDM5 (top), streptavidin (middle), and Tubulin (bottom) are shown. B) Heatmap of 87 KDM5 high confidence interactors (HCIs) and their relative enrichment in TurboID:KDM5^JmjC*^ and TurboID:KDM5^L854F^. C) Volcano plot showing the relative protein abundance of KDM5 HCIs from IP-MS of TurboID:KDM5^JmjC*^ compared to TurboID:KDM5^WT^. Black dots indicate p-value < 0.05, Log2FC ^+^/-1. D) Volcano plot showing the relative protein abundance of KDM5 HCIs from IP-MS of TurboID:KDM5^L854F^ compared to TurboID:KDM5^WT^. Purple dots indicate p-value < 0.05, Log2FC ^+^/-1. E) Venn diagram showing common differential interactors for KDM5^L854F^ and KDM5^JmjC*^. F) GO analysis of HCI proteins that show significantly altered proximity using TurboID:KDM5^L854F^. G) Using STRING-DB and GO-enrichment categories we manually organized proteins with reduced proximity based on complex and biological theme. Colored rings symbolize downregulation in that genotype (black = TurboID:KDM5^JmjC*^, purple = TurboID:KDM5^L854F^). Line width symbolizes degree of connectivity and line darkness indicates confidence in physical interaction.

Biotinylated proteins were purified from adult heads expressing TurboID:KDM5^WT^, TurboID:KDM5^JmjC*^, or TurboID:KDM5^L854F^ and subjected to LC-MS/MS analyses. We constrained our analysis to the 87 high-confidence KDM5 interactors and controlled for changes driven by reduced mRNA expression using our RNA-seq data, although this did not exclude any proteins (log2(FC) +/-1 and p-value<0.05 cutoff; Fig. 4B; SFig 3B-C). TurboID:KDM5^L854F^ showed decreased proximity to 37 proteins, while only seven were reduced using TurboID:KDM5^JmjC*^, two of which were identified in both datasets (FDR>0.05; Fig. 4C-F; Table S2). These data suggest that although the JmjC domain may contribute to the stability of some interactions, a key role for the C5HC2 zinc finger that is altered in KDM5^L854F^ may be to mediate protein-protein interactions. Manual and GO-assisted curation of the proteins showing reduced proximity to TurboID:KDM5^L854F^ revealed an enrichment for proteins involved in chromatin organization and remodeling, including components of the BAF/Brahma and NURF complexes and insulators (Fig. 4G).[36,37] Similar to the H3K4me3 data, this reinforces the notion that while KDM5^JmjC*^ and KDM5^L854F^ cause similar changes to transcription, many molecular features of these variant proteins are distinct.

Proteins such as BEAF-32, Putzig, and nuclear Lamin are insulators that show reduced proximity to KDM5^L854F^ and are modifiers of position effect variegation (PEV) using *In(1)w^m4^*.[38–40] This chromosomal inversion places the *white* gene in proximity to pericentromeric heterochromatin, leading to adult eyes with a mottled red appearance that correlates with the extent to which the *white* gene is expressed or repressed (Fig 5A). [41] Heterozygosity for the null *Kdm5^Δ^*allele enhanced PEV, consistent with prior data using a hypomorphic *P* element allele, as did two alleles of the insulator *Chro* that was identified in our TurboID data (Fig. 5B, C) [42,43]. Similar to *Kdm5^Δ^, Kdm5^L854F^* dominantly enhanced PEV, while *Kdm5^JmjC*^* had no effect (Fig. 5D, E). KDM5 therefore alters PEV in a demethylase-independent fashion and in a manner that correlates with reduced interactions with insulator proteins. PEV can be altered through the spread of heterochromatin that correlates with altered chromatin accessibility.[44] To examine this, we carried out ATAC-Seq comparing adult brains from *Kdm5^L854F^* and *Kdm5^JmjC*^* to *Kdm5^WT^*. Meta-TSS analysis showed a slight increase in signal in *Kdm5^L854F^* and *Kdm5^JmjC*^*, although quantifying these changes revealed few significant differences that did not correlate with altered gene expression (log2FC>+/-1, p-value<0.01; Fig. 5F-H; SFig.4A-I). These data suggest that there are not dramatic changes to overall heterochromatin distribution or chromatin accessibility across the whole brain in either *Kdm5^L854F^* or *Kdm5^JmjC*^*. This is therefore unlikely to account for the observed effect of *Kdm5^L854F^*on PEV, or the shared altered transcriptional programs in *Kdm5^JmjC*^*or *Kdm5^L854F^*.

**Figure 5:**
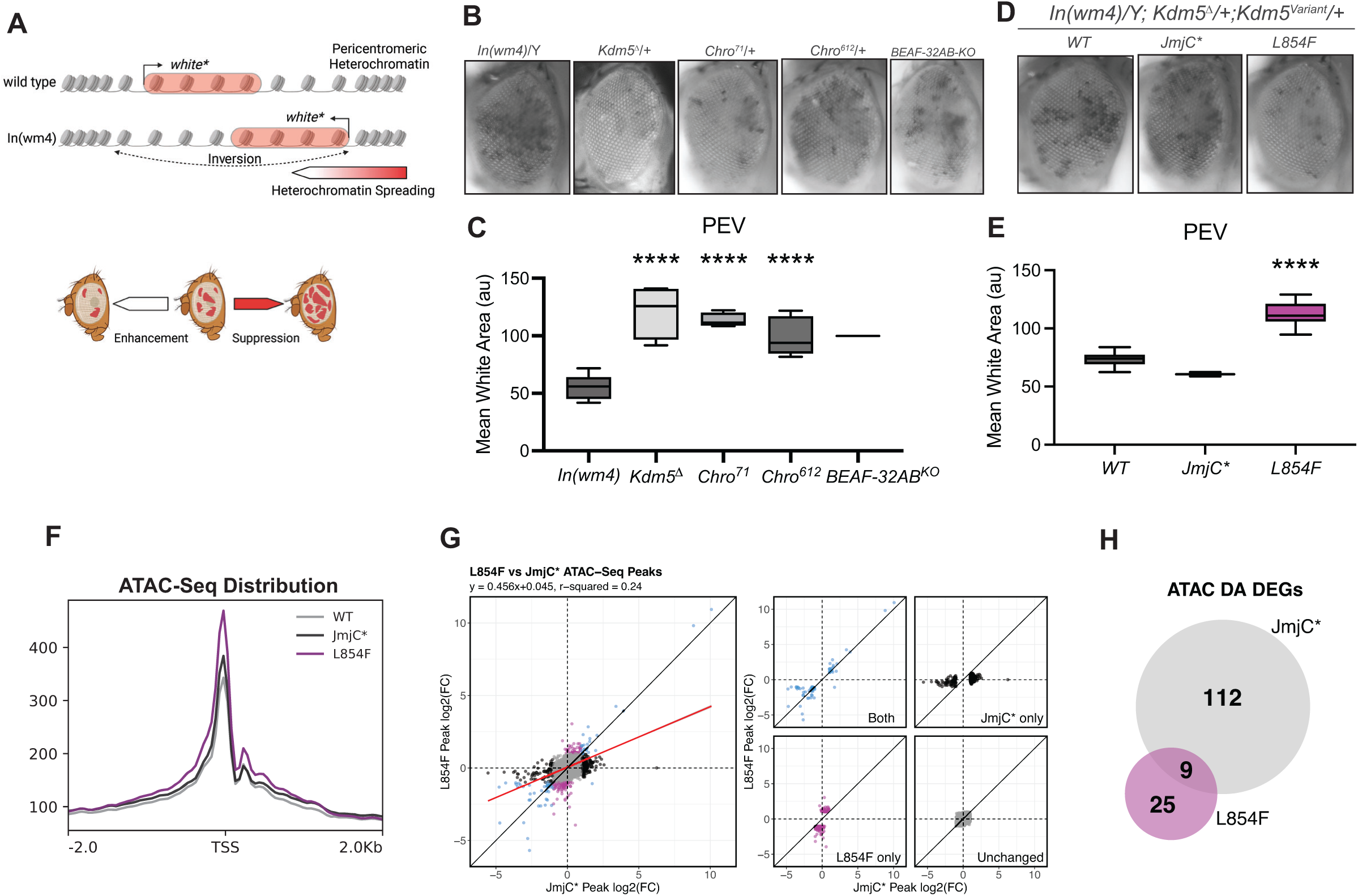
*Kdm5^L854F^* but not *Kdm5^JmjC*^* enhances position-effect variegation. A) Schematic of the *In(1)w^m4h^*inversion allele that places the white gene adjacent to pericentric heterochromatin and is used in the position effect variegation (PEV) assay. B) Representative pictures of adult flies showing PEV seen from crossing *In(1)w^m4h^* to *w^1118^* (control), and enhancement by *Kdm5^Δ^*, *Chro^71^* and *Chro^612^*. C) Quantification of PEV assay by assessing the amount of each eye that has white pigment. D) Representative eye pictures of PEV assays in *In(1)w^m4h^* animals heterozygous for *Kdm5^JmjC*^* or *Kdm5^L854F^*. E) Quantification of PEV assay in ID variants. F) Metaplot showing ATAC-seq surrounding the TSS from *Kdm5^WT^*, *Kdm5^JmjC*^* and *Kdm5^L854F^* animals. G) Scatterplot comparing log2(FC) of ATAC-seq data of *Kdm5^JmjC*^*and *Kdm5^L854F^*. Blue indicates differential accessibility in *Kdm5^JmjC*^* and *Kdm5^L854F^*, purple indicates *Kdm5^L854F^* only, black *Kdm5^JmjC*^* only and grey unchanged in both genotypes. H) Venn diagram showing overlap between genes with altered accessibility in ATAC-seq data and altered gene expression in RNA-seq data.

### *Kdm5^JmjC*^* and *Kdm5^L854F^* disrupt genomic architecture

Insulator proteins are necessary for a range of processes related to proper compartmentalization and organization of the genome.[45,46] To look broadly at whether KDM5 is required for appropriate genome architecture, we performed Hi-C on heads from *Kdm5^WT^*, *Kdm5^JmjC*^*, and *Kdm5^L854F^* flies and generated 25kb resolution contact maps (Fig 6A-C). Based on the effects observed on PEV, we looked for effects on the organization of topologically associating domain (TAD) compartmentalization within the nucleus by examining A and B compartments that are associated with active and inactive chromatin, respectively.[47–49] *Kdm5^JmjC*^* showed A and B compartment proportions that were similar to *Kdm5^WT^* that reflected a small amount of switching from A to B and B to A at similarly low frequencies (2% and 2.5%, respectively; Fig. 6D, E). In contrast, *Kdm5^L854F^* showed a slight bias toward compartment B (55.4%) with A to B compartment switching more than B to A (3,3% and 1.3% respectively), suggesting that some parts of the genome are more heterochromatic (Fig. 6F). Integrating TaDa binding with Hi-C data revealed an enrichment for KDM5 at TAD boundaries (TADb; Fig. 6G). Moreover, TADbs correlated with altered transcription, with the TSSs of downregulated genes being significantly closer to TADbs than upregulated or unchanged genes in *Kdm5^JmjC*^* and *Kdm5^L854F^* (Fig. 6H, I, Table S1). Examining data from *Kdm5^WT^* identified a total of 6728 loops and revealed an enrichment for KDM5 TaDa signal at loop anchors (Fig. 6J). KDM5 also regulates genes present at loop anchors, many of which were similarly differentially expressed in *Kdm5^JmjC*^* and *Kdm5^L854F^*(68% overlap; Fig. 6K). Anchor-associated genes that showed significant dysregulation were enriched for GO terms associated with synapse organization and cytoplasmic translation, which is a key characteristic feature across all ID-variant strains (Fig. 6L). This is also consistent with previous observations that loop anchors often occur near housekeeping genes such as ribosomal protein genes.[50–53] These data suggest that KDM5 may have role at loop anchors and that both KDM5^JmjC*^ and KDM5^L854F^ commonly disrupt this activity.

**Figure 6:**
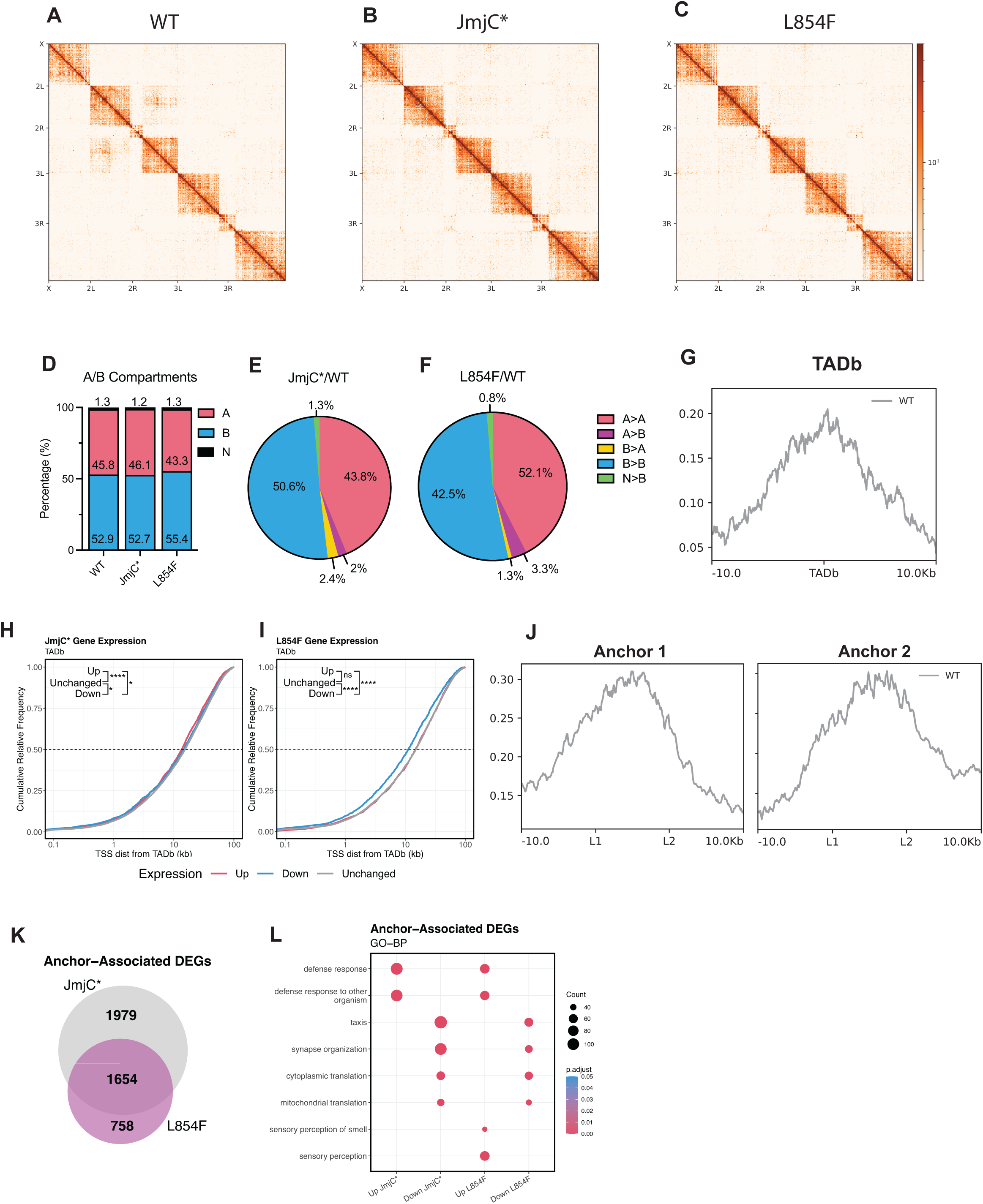
KDM5 affects genome architecture. A) Hi-C contact map (25kb) from *Kdm5^WT^* brains. B) Hi-C contact map (25kb) from *Kdm5^JmjC*^* brains. C) Hi-C contact map (25kb) from *Kdm5^L854F^* brains. D) Quantitation of compartments A and B total amounts in *Kdm5^WT^*, *Kdm5^JmjC*^* and *Kdm5^L854F^*. E) Alterations in A/B compartments comparing *Kdm5^JmjC*^* to *Kdm5^WT^*. F) Alterations in A/B compartments comparing *Kdm5^L854F^* to *Kdm5^WT^*. G) Metaplot showing enrichment of KDM5^WT^ at TAD boundaries (TADb). H) Distance from KDM5-bound TADb and altered gene expression in *Kdm5^JmjC*^*. I) Distance from KDM5-bound TADb and altered gene expression in *Kdm5^L854F^*. J) Meta-analysis of TaDa data at loops anchors called in *Kdm5^WT^*. K) Venn diagrams showing DEGs from *Kdm5^JmjC*^* and *Kdm5^L854F^* of genes present at loop anchors called in *Kdm5^WT^*. L) GO-BP analysis of anchor-associated DEGs in *Kdm5^JmjC*^* and *Kdm5^L854F^*.

To further explore the role of KDM5 in genomic organization, we compared Hi-C data from *Kdm5^JmjC*^* and *Kdm5^L854F^*to *Kdm5^WT^*. This revealed a strikingly similar overall pattern of changes in the two genotypes (Fig. 7A, B). Local TADs were not generally disrupted in *Kdm5^JmjC*^* or *Kdm5^L854F^* brains, as indicated by the white along diagonal in the contact maps. The most obvious changes in *Kdm5^JmjC*^* and *Kdm5^L854F^* were related to chromatin looping. For example, on chromosome II in both *Kdm5^JmjC*^*and *Kdm5^L854F^* three large loops, several local loops and an intra-chromosomal loop between the chromosomal arms were all decreased (Fig. 7A, B, SFig. 5A-D). 10kb binned contact maps showed an enrichment for KDM5 TaDa signal at the anchors of these altered genomic loops (SFig 5A-D). Differential loop calling analyses revealed that *Kdm5^JmjC*^* gained 113 and lost 96 loops, while *Kdm5^L854F^*gained 232 and lost 61 loops (Fig. 7C). A zoomed in view of chromosome 2L in both *Kdm5^JmjC*^* and *Kdm5^L854F^* show an overall similar pattern in the contact matrix, however there are more gained loops throughout the chromosome in *Kdm5^L854F^* compared to *Kdm5^JmjC*^* (Fig. 7D, E). Aggregate Peak Analysis (APA) suggested that lost loops were common between genotypes but that gained loops were distinct (Fig 7F, G). These data are consistent with the variants *Kdm5^JmjC*^* and *Kdm5^L854F^* disrupting unique and shared activities related to the formation or maintenance of chromatin loops.

**Figure 7:**
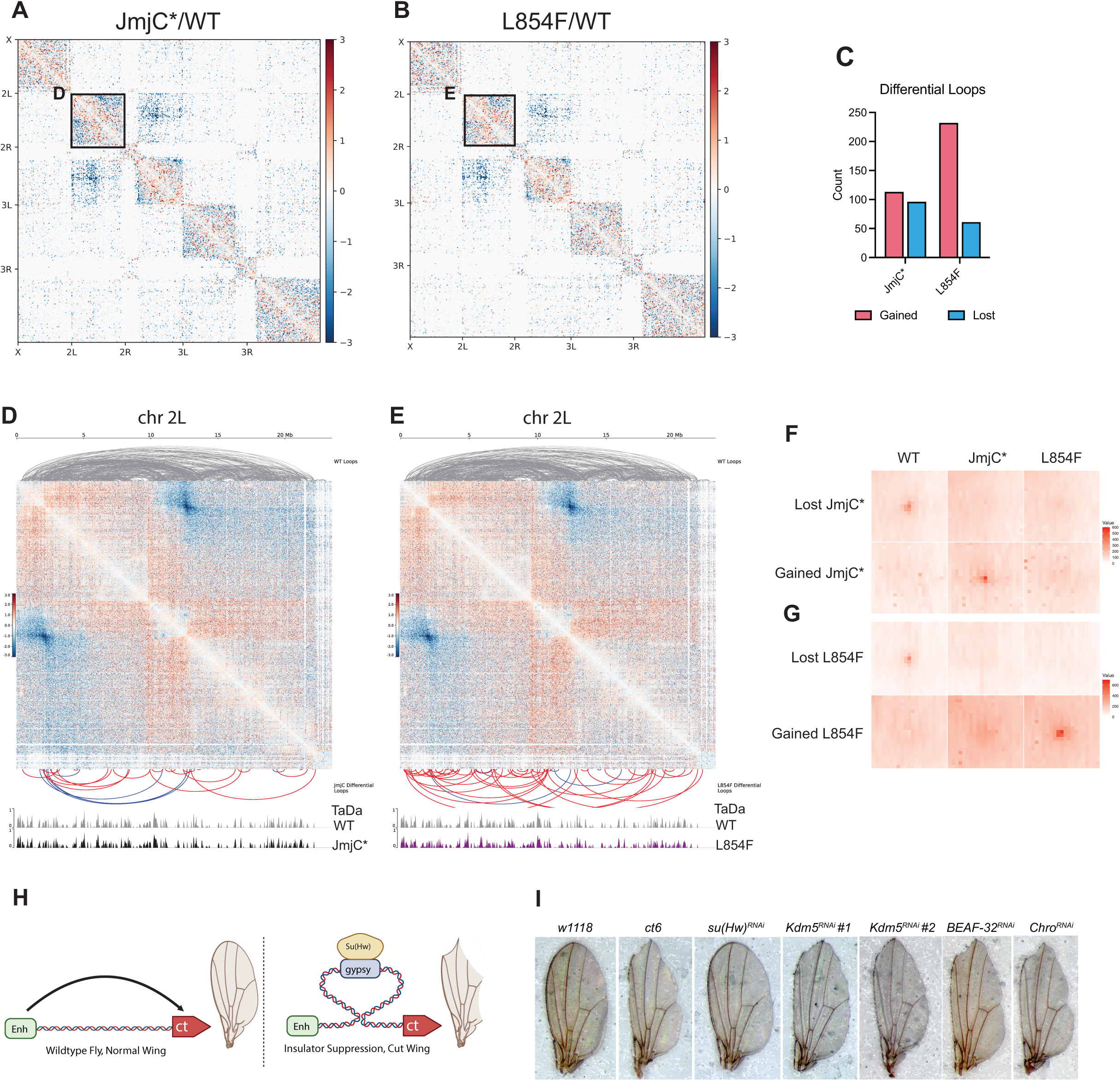
KDM5^JmjC*^ and KDM5^L854F^ alter loops. A) Hi-C contact map (25kb) comparing *Kdm5^JmjC*^* to *Kdm5^WT^* B) Hi-C contact map (25kb) comparing *Kdm5^L854F^* brain to *Kdm5^WT^* C) Bar graph of differential loops called per genotype. D) Hi-C contact map (10kb) of chr2L comparing *Kdm5^JmjC*^* brain to *Kdm5^WT^* zoomed in at black box indicated in A. Top gray loops represent loops called in *Kdm5^WT^*. Bottom panel are gained (red) and lost (blue) differential loops. Genome tracks represent KDM5 TaDa signal. E) Hi-C contact map (10kb) of chr2L comparing *Kdm5^JmjC*^* brain to *Kdm5^WT^* zoomed in at black box indicated in A. Top gray loops represent loops called in *Kdm5^WT^*. Bottom panel are gained (red) and lost (blue) differential loops. Genome tracks represent KDM5 TaDa signal. F) APA analysis of gained and lost loops in *Kdm5^JmjC*^*. G) APA analysis of gained and lost loops in *Kdm5^L854F^*. H) Schematic of the *cut^6^* allele and the effects of the mdg4 retrotransposon that interferes with enhancer-promoter interactions. I) Representative images wings from the *cut^6^* flies using the wing disc driver nubbin-Gal4 to knockdown *su(Hw)* (positive control), *Kdm5*, *BEAF-32* or *Chro*.

To further examine the functional role of KDM5 in looping, we exploited a well-characterized genetic assay using the *cut^6^* allele.[54–56] The notched wing phenotype of this allele is caused by the insertion of mdg4 (formerly known as gypsy) retrotransposon that blocks enhancer-promoter interactions and causes loss of wing margin expression of *cut* during development (Fig. 7H).[54] RNAi-mediated knock down of *Kdm5* using two independent sh-RNAi lines during wing development suppressed the *cut^6^*wing notching phenotype (Fig. 7I). Interestingly, the *cut^6^* wing phenotype was not altered by knocking down other insulator proteins known to alter PEV that showed changes in the KDM5^L854F^ TurboID data, including *BEAF-32* and *Chro* (Fig. 7I). This reinforces the notion that the effects of the C5HC2 domain of KDM5 linked to PEV are likely distinct from those needed for looping. KDM5 therefore functions as an insulator in part by maintaining loop architecture.

## Discussion

Here we focus on understanding the mechanisms by which KDM5 functions to regulate gene expression in the brain using the ID-associated allele *Kdm5^L854F^*. Prior studies have shown that this allele displayed similar adult behavioral phenotypes and gene expression changes to the demethylase inactive *Kdm5^JmjC*^* allele.[9–11] Despite these similarities, examining bulk levels of H3K4me3 showed that *Kdm5^L854F^* did not recapitulate the increase in this chromatin mark seen in *Kdm5^JmjC*^*. Consistent with the primary deficit of the KDM5^L854F^ protein being unrelated to its enzymatic activity, we show that the distribution of H3K4me3 at promoters was much more similar to wild type than to the broadened peaks seen in *Kdm5^JmjC*^*brains. Rather, a key deficit of the KDM5^L854F^ is an altered ability to properly interact with other proteins, particularly those that function as insulators. This correlated with the ability of *Kdm5^L854F^*, and not *Kdm5^JmjC*^*, to enhance PEV, indicating a possible demethylase-independent role in heterochromatin formation for KDM5. In contrast, both variants showed similarly altered activities at loop anchors, including the altering the expression of genes such as those necessary for translation. In agreement with KDM5 possessing activities critical to looping, *Kdm5* knockdown suppressed mdg4-mediated insulation of enhancer-promoter activity. Combined, these data show that KDM5 has insulator activity and that two distinct regions, the catalytic JmjC domain, and the C5HC2 zinc finger, have distinct and overlapping functions, both of which are likely to be important for proper brain development and function.

The pathogenic *Kdm5^L854F^* variant affects the C5HC2 zinc finger motif of KDM5 that is immediately adjacent to the JmjC domain, and is considered to be part of the catalytic cassette.[57] In keeping with this, *in vitro* studies of KDM5C^L731F^, the human equivalent of KDM5^L854F^, showed a 50% decrease in activity toward a histone peptide substrate.[27] However, *in vivo*, KDM5^L854F^ results in only mild effects on H3K4me3, possibly reflecting the more robust activity of this variant protein as part of physiologically relevant complexes and/or against nucleosomal substrates. While we cannot exclude the possibility that minor changes to KDM5^L854F^ demethylase activity could contribute to phenotypes, it is more likely that phenotypes are being driven by altered protein-protein interactions. Interestingly, the C5HC2 region appears to be a hotspot for ID-associated variants in KDM5C, with at least six other missense variants occurring in different amino acids within this small domain.[1] We have generated *Drosophila* models for two of these, allowing us to determine whether they show similar changes in proximity to interactors as KDM5^L854F^, and whether these are distinct from variants that impact other domains of KDM5.[9]

Similar to proximity-labeling studies, the extent to which other ID-alleles affect PEV will also provide insights into the role of KDM5 in this process. The enhancement of PEV observed by heterozygosity for *Kdm5^Δ^* and *Kdm5^L854F^*, but not *Kdm5^JmjC*^*, suggests a demethylase-independent role for KDM5 in maintaining permissive chromatin environments. Although we were unable to observe differences in accessibility using ATAC-seq, possibly due to our use of whole brains, we did find that *Kdm5^L854F^* and not *Kdm5^JmjC*^* caused a mild shift from compartment A to B in our Hi-C data. How KDM5, and specifically its C5HC2 domain, functions to mediate this role remains to be determined. One possible mechanism is through altered interactions with BEAF-32 and other insulators that behave as enhancers of PEV and show reduced proximity to KDM5^L854F^ and not KDM5^JmjC*^.[38–40] KDM5-insulator interactions could be required for the proper localization of heterochromatin within the nucleus or in forming a boundary that limits the spread of heterochromatin. It is also possible that KDM5 more directly impacts the distribution or recognition of H3K9 methylation, a chromatin mark that characterizes pericentric constitutive heterochromatin. This is based on the observation that *Kdm5* genetically interacts with the H3K9 methyltransferase G9a, although the underlying mechanism for this remains unknown.[58] Whatever the cause, the observation that *Kdm5^JmjC*^* and *Kdm5^L854F^* show distinct behaviors with respect to PEV suggests that this C5HC2-mediated function of KDM5 is distinct from its role in influencing genomic architecture.

The deficits of KDM5^JmjC*^ and KDM5^L854F^ variant proteins converge on a shared phenotype related to activities at loop anchors. This raises the key question of how variants that show distinct effects on enzymatic activity and protein-protein interactions cause similar effects at these regions. One possibility is that both the JmjC domain-encoded enzymatic activity and C5HC2 zinc finger-mediated protein-protein interactions are necessary for this activity. This is an unlikely scenario, particular as the many genes, including ribosomal protein genes that often occur near loop anchor regions, did not show significant changes to H3K4me3 in *Kdm5^JmjC*^*. Instead, it is perhaps more likely that altered complex formation drives the loop phenotypes of both alleles. While CTCF and other zinc finger proteins work with Cohesin to influence loop and TAD formation in mammalian cells, these proteins play more limited roles in *Drosophila*.[59] One feature that is conserved across species is the presence of housekeeping genes, including ribosomal protein genes, near loop anchor regions.[50,52,60,61] One model that would be compatible with our data is that KDM5-regulated transcription of genes near loop anchors is important for the integrity of the loop as a whole. Consistent with this, while there was limited overlap observed in the TurboID data using KDM5^L854F^ and KDM5^JmjC*^, both variants showed reduced interactions with members of the Mediator complex.[35,62] Mediator plays roles in RNA Pol II transcriptional initiation, elongation, pausing and reinitiation.[63–65] Similarly, KDM5 demethylase activity has been shown to aid in RNA Pol II initiation and pause release.[15,31] KDM5^L854F^ has reduced proximity to MED14, and KDM5^JmjC*^ to MED17 using a p<0.05 cutoff, although this expands to MED1, MED14, and MED17 for both mutants using p<0.14. Interactions between KDM5 and MED14, MED17, and/or MED1, which are adjacent each other in the assembled Mediator complex, may be particularly important at genes near loop anchors, and this could disrupt looping. This could be a key pathomechanism for the altered cognition observed in *Kdm5^JmjC*^* and *Kdm5^L854F^*, as components of the Mediator complex are implicated in a range of neurological disorders.[66] Of particular note is the observation that *MED17* alleles have been associated with phenotypes similar to those seen in individuals with variants in *KDM5A*, *KDM5B* or *KDM5C* such as intellectual disability, seizures, and autism.[67,68] Defining where components of Mediator are bound in the genome of neurons, and how this is altered in ID variant strains, will be paramount to uncovering this mechanism.

The downstream consequences of KDM5-associated activity at anchor regions on local looping remain to be further explored, as the resolution of Hi-C data do not allow accurate identification of the specific genes affected. In this regard, it is notable that although *Kdm5^JmjC*^* and *Kdm5^L854F^* show similar transcriptional changes at loop anchor regions, the changes to chromatin looping were not the same. Indeed, *Kdm5^L854F^* showed many more changes, particularly gained loops, than *Kdm5^JmjC*^*. Looking to the future, higher resolution data are needed, as are additional 3D-genome organization studies using other ID-alleles in *Drosophila* and in mammalian cells. Combined, these studies will define how KDM5 family proteins regulate gene expression programs in the brain and their link to Claes-Jensen syndrome.

## Materials and Methods

### Resource availability

Further information and requests for reagents should be directed to Julie Secombe.

### Fly strains and care

Fly food (per liter) contained 80g malt extract, 65g cornmeal, 22g molasses, 18g yeast, 9g agar, 2.3g methyl para-benzoic acid and 6.35ml propionic acid per liter. Flies were kept at 25°C with a 12-hour light/dark cycle and 50% humidity. All flies were collected or harvested 1-5 days after eclosion from 9am-12pm. *Kdm5*^Δ^ (null allele; previously *Kdm5^140^*), *Kdm5^A224T^*, *Kdm5^JmjC*^*, *Kdm5^L854F^*, are published.[9,18,19] UASp-TurboID:*Kdm5* constructs were generated by site-directed mutagenesis of published UASp-TurboID:*Kdm5*.[35] These transgenes were inserted into attP at 86F. For PEV analyses, the *white* gene and RFP cassette was removed. UAS-Dam:*Kdm5*-ID variant constructs were generated by site-directed mutagenesis of published UAS-Dam:*Kdm5*.[19] All Dam fusion transgenes were inserted into attP2 on chromosome III. All transgenesis were carried out at Best Gene. The following fly stocks were obtained from Bloomington *Drosophila* Stock Center: *w^1118^* (RRID:BDSC_5905), Ubi-Gal4 (RRID:BDSC_32551), *In(1)w^m4h^* (RRID: BDSC_76618), *Chro^71^* (RRID:BDSC_93149), *Chro^612^* (RRID:BDSC_93148), *nubbin* (nub)-Gal4 (RRID:BDSC_862108). RNAi lines: *Kdm5 (#1)* (RRID:BDSC_35706), *Kdm5 (#2)* (RRID:BDSC_28944), *su(Hw)* (RRID: BDSC_33906).

### Antibodies

The following primary antibodies were used: Anti-HA (1:1000, Cell Signaling Technology cat #2367, RRID:AB_10691311), anti-α-Tubulin (1:5000, Cell Signaling Technology cat #3873, RRID:AB_1904178), anti-H3 (Abcam24384), anti-H3K4me3 (Abcam8580). Secondary antibodies: IRDye 680RD donkey anti-mouse IgG (1:5000; LI-COR Biosciences cat# 925-68072, RRID:AB_2814912) and IRDye 800CW donkey anti-rabbit IgG (1:5000; LI-COR Biosciences cat# 926-32213, RRID:AB_621848). Western Blots were scanned and processed using a LI-COR Odyssey Infrared scanner.

### Targeted DamID (TaDa)

Tissue processing was performed as previously described with the following modifications: TaDa was performed in triplicate (*Kdm5^L854F^* in duplicate) for each genotype with 10 heads per sample.[19] KDM5-DamID fusion proteins were expressed under the control of Ubi-Gal4, which was pulsed on for three days during early adulthood. Genotypes were *Kdm5^Δ^*, Ubi-Gal4/*Kdm5^Δ^*, tubulin-Gal80^ts^; UAS-*Kdm5:DamID^WT^*or variant, *gKdm5:HA^WT^ ^or^ ^variant^*. Samples were homogenized in 75μl UltraPure Distilled Water and 20 μl 500 mM EDTA then digested with Proteinase K for 1.5h. DNA extraction was performed using the Zymo Quick-DNA Miniprep Plus Kit (Zymo, D3024). DpnI digestion, PCR adaptor ligation, DpnII digestion, and PCR amplification were performed as described. DNA was sonicated using a Diagenode Bioruptor Pico for 6 cycles (30s on/90s off at 4°C), and DNA fragments were analyzed using an Agilent Bioanalyzer to confirm ∼300 bp fragment size. DamID adaptor removal and DNA cleanup were performed as previously described, and samples were submitted to BGI Genomics for library construction and sequencing. Libraries were prepared at BGI Genomics following a ChIP-seq workflow and sequenced on the NovaSeq S4.

For TaDa analyses, sequencing data were aligned to the *D. melanogaster* dm6 genome and processed using damidseq_pipeline.[69] After converting to bedgraphs via damidseq_pipeline, peaks were called using find_peaks (using the parameters fdr=0.05, min_quant=.8) on the averaged replicates, and genes overlapping peaks identified using ChIPSeeker.[70] We used NOISeq to perform differential analysis with a cutoff of 0.8 as describe in Xu et al.[71]

### Single-Cell Suspension

50 brains were dissected and placed in 1 mL of ice-cold 1X PBS. Brains were centrifuged at 4C for 5 minutes at 1000XG. PBS was removed and the cell pellet was resuspended with 150 uL of 100 mg/mL Collagenase, Type I (ThermoFisher, 17018029) and incubated at 25C on a thermoshaker at 500 rpm for 2 hours. The suspension was vigorously pipetted up and down 20 times every 10-15 minutes to facilitate digestion. After incubation, suspensions were centrifuged again centrifuged at 4C for 5 minutes at 1000XG and resuspended in PBS + 10% DMSO. 10 uL of the suspension was used for hemocytometer. 50 brains yielded about 1.5 million cells. The remaining 100 uL was slow-frozen with a Mr. Frosty Freezing Container (ThermoFisher, 5100-0001) at –80C.

### CUT & RUN

CUT & RUN was performed with Epicypher CUTANA Kit (Epicypher, 14-1048) and Epicypher Library Prep Kit (Epicypher, 14-1001) as directed using generated single cell suspension. Antibodies against H3K4me3 (Epicypher Cat# 13-0041, RRID:AB_3076423), and IgG (Epicypher Cat# 13-0042, RRID:AB_2923178). Samples were sequenced on the Illumina MiSeq Platform. All samples were normalized to spike-in *E. coli* DNA.

### ATAC-Seq

ATAC-Seq was performed using the ATAC-Seq Kit from Active Motif (Active Motif, 53150) using manufacturer’s instructions. Samples were sequenced on the Illumina MiSeq Platform.

### Peak Alignment and Analysis

Raw reads quality controlled and trimmed using rfastp (v 1.6.0) and were aligned using Rsubread (2.10.5) to the dm6 *Drosophila melanogaster* genome assembly.[72,73] BAM files were normalized through dividing number of E.coli reads in each sample by the lowest number of E. coli reads across samples. Samtools (v 1.14) was then used to scale BAM files by this normalization factor using (samtools view –s (normalization factor)).[74] BigWigs were generated using deeptools (v 3.5.1) bamCoverage function.[75] Profiles and heatmaps were generating using deeptools computeMatrix and plotProfile functions. Peaks were called using Genrich(v 0.6.1) in triplicate, DiffBind (v3.6.1) was used to perform differential methylation analyses.[76,77] Reads in each peak were counted using dba.count(summits = FALSE). ChIPSeeker (v1.32.0) was used to perform peak annotation and clusterProfiler(v4.4.4) to perform Gene Ontology Analyses.[70,78]

### TurboID-mediated Biotinylated Protein Enrichment

2-5 day-old flies were flash frozen in liquid nitrogen and decapitated. Using 10 heads per replicate, samples were homogenized in 250 μL RIPA Buffer (Thermofisher 89901) supplemented with Halt™ Protease Inhibitor Cocktail (Thermofisher, 78430) and centrifuged at 4 °C for 10 minutes at 15,000XG to remove debris. 100 μL of Pierce™ Streptavidin Magnetic Beads (Thermofisher, 88817) were washed twice with RIPA and the cleared lysate was added. The lysate-bead mixture was incubated with rotation at 4 °C overnight. The next day the lysate was discarded, and beads were washed twice with RIPA, once with 1M KCl, once with 0.1 M Na_2_HCO_3_, once with 1 M Urea in 10 mM Tris pH 8.0, and twice again with RIPA. For Western Blot analyses all RIPA was removed and biotinylated proteins were eluted with 4X NuPAGE™ LDS Sample Buffer (Invitrogen, NP0007) supplemented with 2 mM biotin and 20 mM DTT.

### On-bead protein digestion

Proteins were digested directly on streptavidin beads. 5 mM DTT and 50 mM ammonium bicarbonate (pH = 8) were added to the solution and left on the bench for about 1 hour for disulfide bond reduction. Samples were then alkylated with 20 mM iodoacetamide in the dark for 30 minutes. Afterward, 500 ng of trypsin was added to the samples, which were digested at 37 °C for 18 h. The peptide solution was dried in a vacuum centrifuge.

### Sample desalting

Prior to mass spectrometry analysis, samples were desalted using a 96-well plate filter (Orochem) packed with 1 mg of Oasis HLB C-18 resin (Waters). Briefly, the samples were resuspended in 100 µl of 0.1% TFA and loaded onto the HLB resin, which was previously equilibrated using 100 µl of the same buffer. After washing with 100 µl of 0.1% TFA, the samples were eluted with a buffer containing 70 µl of 60% acetonitrile and 0.1% TFA and then dried in a vacuum centrifuge.

### LC-MS/MS Acquisition and Analysis

Samples were resuspended in 10 µl of 0.1% TFA and loaded onto a Dionex RSLC Ultimate 300 (Thermo Scientific), coupled online with an Orbitrap Fusion Lumos (Thermo Scientific). Chromatographic separation was performed with a two-column system, consisting of a C-18 trap cartridge (300 µm ID, 5 mm length) and a picofrit analytical column (75 µm ID, 25 cm length) packed in-house with reversed-phase Repro-Sil Pur C18-AQ 3 µm resin. Peptides were separated using a 90 min gradient from 4-30% buffer B (buffer A: 0.1% formic acid, buffer B: 80% acetonitrile + 0.1% formic acid) at a flow rate of 300 nL/min. The mass spectrometer was set to acquire spectra in a data-dependent acquisition (DDA) mode. Briefly, the full MS scan was set to 300-1200 m/z in the orbitrap with a resolution of 120,000 (at 200 m/z) and an AGC target of 5×10e5. MS/MS was performed in the ion trap using the top speed mode (2 secs), an AGC target of 1×10e4 and an HCD collision energy of 35. Raw files were searched using Proteome Discoverer software (v2.4, Thermo Scientific) using SEQUEST search engine and the UniProt database of Drosophila melanogaster. The search for total proteome included variable modification of N-terminal acetylation, and fixed modification of carbamidomethyl cysteine. Trypsin was specified as the digestive enzyme with up to 2 missed cleavages allowed. Mass tolerance was set to 10 pm for precursor ions and 0.2 Da for product ions. Peptide and protein false discovery rate was set to 1%. Following the search, data was processed as described by Aguilan et al.(98). Briefly, proteins were log2 transformed, normalized by the average value of each sample and missing values were imputed using a normal distribution 2 standard deviations lower than the mean. Statistical regulation was assessed using heteroscedastic T-test (if *p*-value < 0.1). Data were assumed to be Gaussian distributed.

### Quantification of In(1)w^m4h^ phenotypes

Pictures of adult heads were taken with a light microscope and converted to black and white using Adobe Photoshop. The magnetic lasso tool was used to trace the eye and the measure tool was used to get Mean Gray Value, which was plotted for one eye per animal.

### Quantification of cut6 wing phenotype

Pictures of 10-15 wings per genotype were taken with a light microscope. A scale of 1-3 was assigned with 3 being a wildtype wing with no defects, and 1 being a fully cut wing. Any restoration of the wing margin was considered 2. Wings were rated blindly.

### Hi-C

We used Arima HiC Kit (A510008) and Arima Library Prep Kit (A303011) using manufacturer’s recommendations. We performed deep sequencing using Illumina NextSeq 2000.Contact maps were created using the Juicer package (v1.6) [79].

### Compartments

Eigenvectors were calculated using JuicerTools as described in Chathoth et al.[48,80] In short, eigenvectors and GC content were calculated for 10 kb bins across each chromosome. The sign of the eigenvectors was corrected for each chromosome such that the correlation was positive. Bins with positive values were assigned as A compartments and negative B compartments.

### Chromatin Loops

Chromatin loops were called with the GPU version of the HiCCUPS tool.[79] Loops were called using a 5-kb resolution, 0.05 FDR, Knight-Ruiz normalization, a window of 10, peak width of 5, thresholds for merging loops of 0.02, 1.5, 1.75, 2, and distance to merge peaks of 20 kb (-k KR –r 2000 –f 0.05 –p 5 –i 10 –t 0.02,1.5,1.75,2 –d 20000) as described in Chathoth et al. 2022.[48] HiCCUPS Diff was used do calculated differential peaks.

### Topologically Associated Domains (TADs)

HiCExplorer (v3.7.5) was used to call TADs at 5 kb, 10kb, and 25 kb resolutions using and merged the hicFindTADs function and merged using the hicMergeDomains function with default settings.[81]

### Plotting

Plotting of genomic regions was done using PyGenomeTracks (v3.8) and HiCExplorer.[81,82]

## Acknowledgements

We thank Hayden Hatch for producing the ID-variant Targeted DamID fly stocks, Owen Marshall for invaluable discussions about TaDa analyses, and members of the Secombe lab, Einstein chromatin club and the Einstein Intellectual and Developmental Disabilities Research Center (IDDRC) for their support and feedback throughout this project. We appreciate the availability of fly strains from the Bloomington *Drosophila* Stock Center (NIH P400D018537) and antibodies from the Developmental Studies Hybridoma bank, created by the NICHD of the NIH and maintained at The University of Iowa. We also thank the Analytical Imaging (shared instrument grant 1S10OD023591), Flow Cytometry and Genomics core facilities at Einstein and the Einstein Cancer Center Support Grant (P30 CA013330).

## Funding

National Institutes of Health grant T32GM149364 (MY)

National Institutes of Health grant F31GM146347 (MY)

National Institutes of Health grant R01GM1112783 (JS)

National Institutes of Health grant R01GM150189 (JS)

National Institutes of Health grant F31GM150194 (MC)

National Institutes of Health grant T32GM145438 (RDK, MC)

National Institutes of Health grant 1-S10-OD030286 (S.S)

Irma T. Hirschl Trust (JS)

American Federation for Aging Research (AFAR) Sagol Networks GerOmics award (S.S)

## Author Contributions

Conceptualization, MY, JS

Methodology MY, RDK, MC, SS

Investigation, MY, RDK, MC, SS, JS

Writing – original draft: MY and JS

Writing – reviewing and editing MY, MC, RDK, MC, SS, JS

## Data and materials availability

All reagents generated in this study are available upon request.

*Kdm5^JmjC*^* and *Kdm5^L854F^* RNA-seq data available at GEO:GSE100578 and GEO:GSE245380.[9,11]

TurboID data are publicly available through ProteomeXchange (project PXD057637). Data is available under BioProject reference: PRJNA1188621 and GEO as follows: H3K4me3 CUT&RUN (GSE282535), Targeted DamID (GSE282537), ATAC-Seq (GSE282534), and HiC (GSE282536).

## Supplemental Figure Legends

**Supplemental Figure 1.**
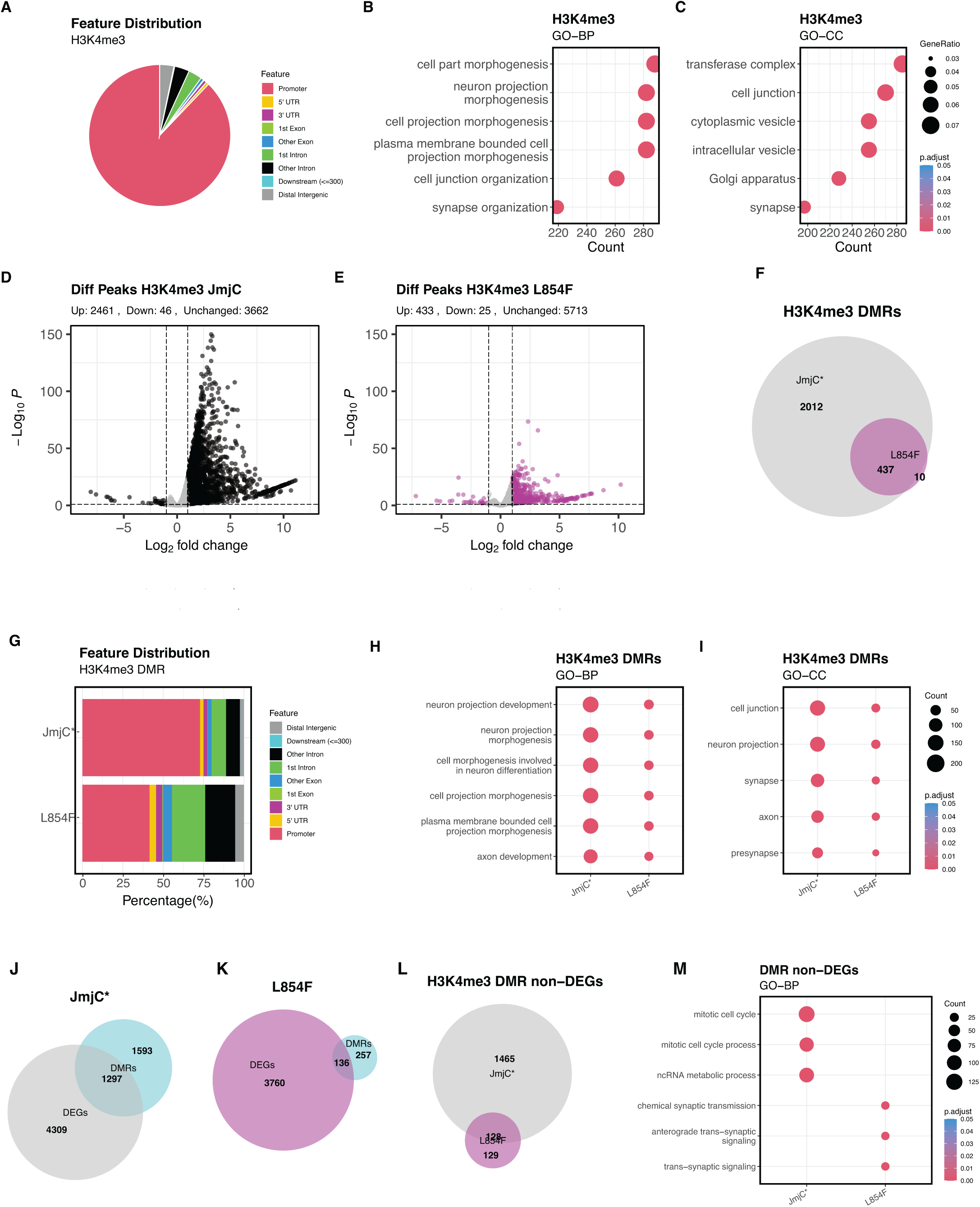
A) Pie chart of H3K4me3 peak annotations. B) GO-BP of H3K4me3 promoter-proximal peaks. C) GO-CC of H3K4me3 promoter-proximal peaks. D) Volcano plot showing changes to H3K4me3 in *Kdm5^JmjC*^*. Black dots indicate statistically significant changes to H3K4me3 peaks. E) Volcano plot showing changes to H3K4me3 in *Kdm5^L854F^*. Purple dots indicate statistically significant changes to H3K4me3 peaks. F) Venn diagram showing the overlap in differential regions of H3K4me3 called in *Kdm5^JmjC*^* and *Kdm5^L854F^*. G) Bar plot of differentially methylated regions (DMR) peak annotations in *Kdm5^JmjC*^* and *Kdm5^L843F^*. H) GO-BP analysis of DMRs from *Kdm5^JmjC*^* and *Kdm5^L843F^*. I) GO-CC analysis of DMRs from *Kdm5^JmjC*^* and *Kdm5^L843F^*. J) Venn diagram of the overlap between genes that show different levels of H3K4me3 (DMRs) and altered transcription (DEGs) in *Kdm5^JmjC*^*. K) Venn diagram of the overlap between genes that show different levels of H3K4me3 (DMRs) and altered transcription (DEGs) in *Kdm5^L854F^*. L) Venn Diagram showing overlap between genes with DMR but not altered expression (non-DEGs). M) GO-BP of genes that were DMR and non-DEGs.

**Supplemental Figure 2.**
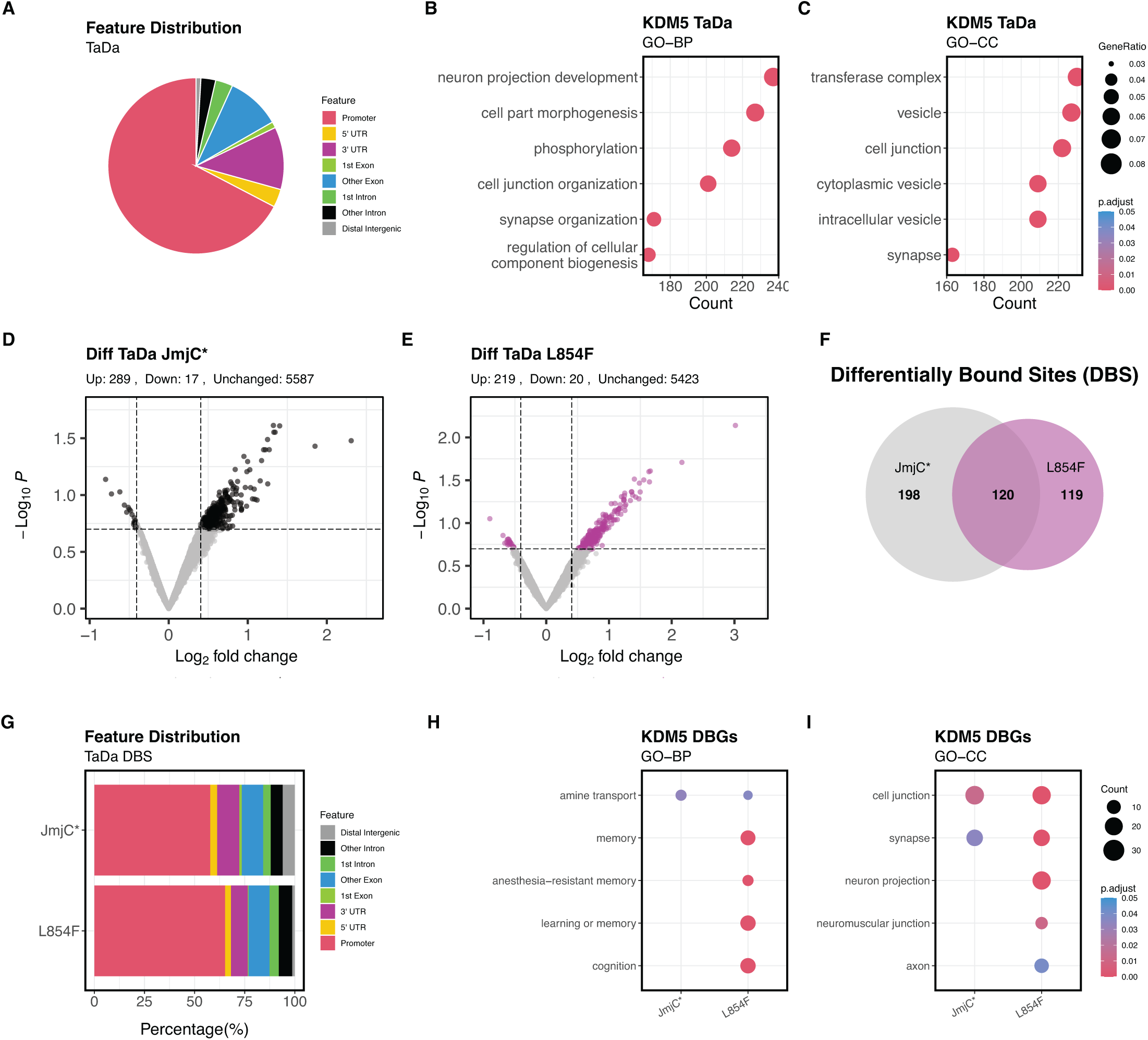
A) Pie chart showing distribution of wild-type KDM5 TaDa peak annotations. B) GO-BP of TaDa promoter-proximal peaks for wild-type KDM5. C) GO-CC of TaDa promoter-proximal peaks wild-type KDM5. D) Volcano plot showing the changes observed in KDM5^JmjC*^ TaDa data. Dots in black indicate significantly altered. E) Volcano plot showing the changes observed in KDM5^L854F^ TaDa data. Dots in purple indicate significantly altered. F) Venn diagram showing the shared differentially bound sites (DBS) in the KDM5^JmjC*^ and KDM5^L854F^ TaDa datasets. G) Bar plot of DBS peak annotations for KDM5^JmjC*^ and KDM5^L854F^ TaDa datasets. H) GO-BP analyses of differentially bound genes (DBGs) KDM5^JmjC*^ and KDM5^L854F^ TaDa datasets. I) GO-CC analyses of differentially bound genes (DBGs) KDM5^JmjC*^ and KDM5^L854F^ TaDa datasets.

**Supplemental Figure 3.**
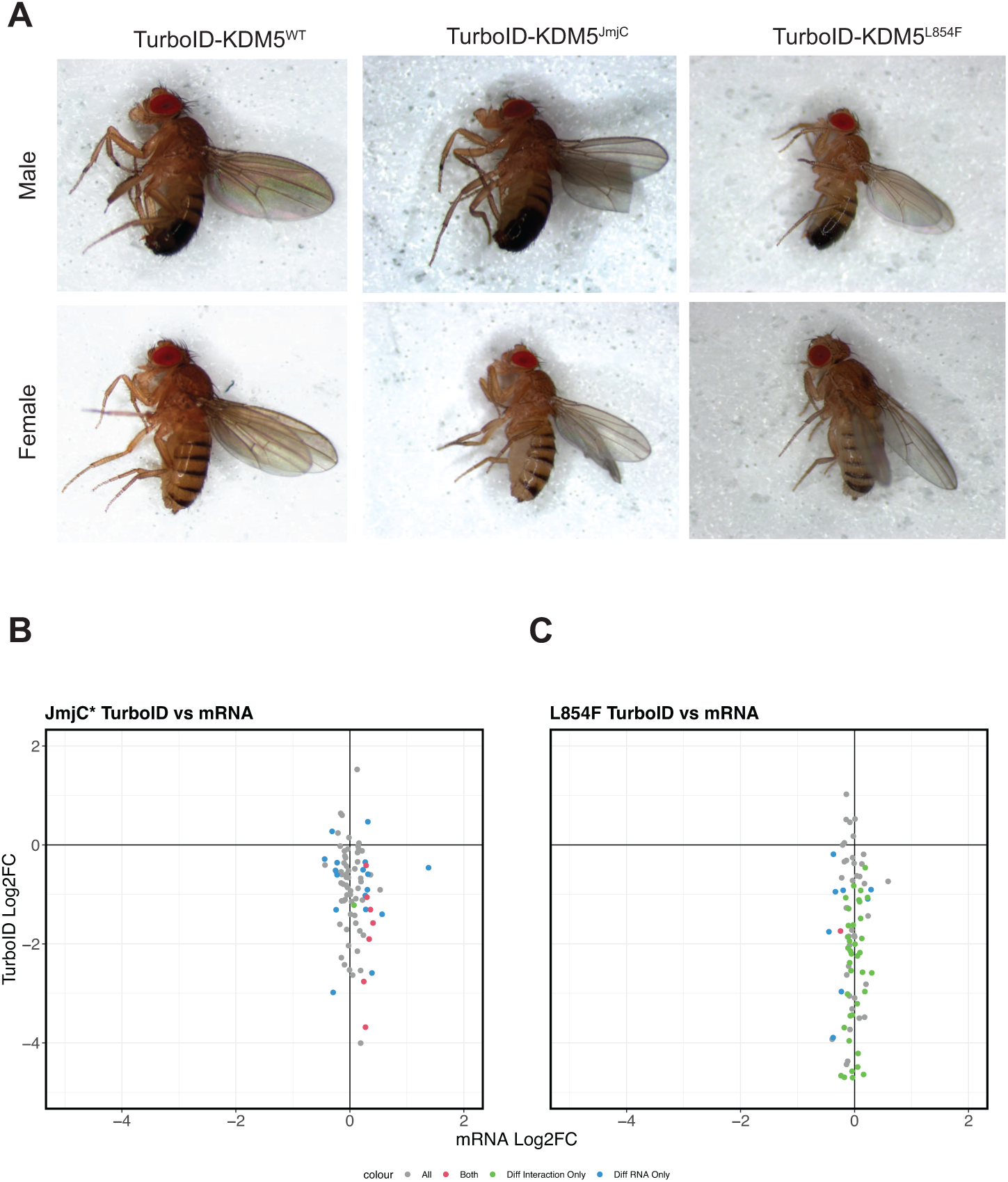
A) Representative pictures of adult male and female flies showing rescue of lethality by the Turbo:KDM5 fusion proteins. Genotypes are *Kdm5^Δ^*, Ubi-Gal4/*Kdm5^Δ^*; UAS-TurboID:*Kdm5^WT^* or *Kdm5^variant^*. B) Scatterplot showing log2 fold change in mRNA seen in *Kdm5^JmjC*^* compared to the relative abundance of proteins identified by TurboID:KDM5^JmjC*^. C) Scatterplot showing log2 fold change in mRNA seen in *Kdm5^L854F^* compared to the relative abundance of proteins identified by TurboID:KDM5^L854F^.

**Supplemental Figure 4.**
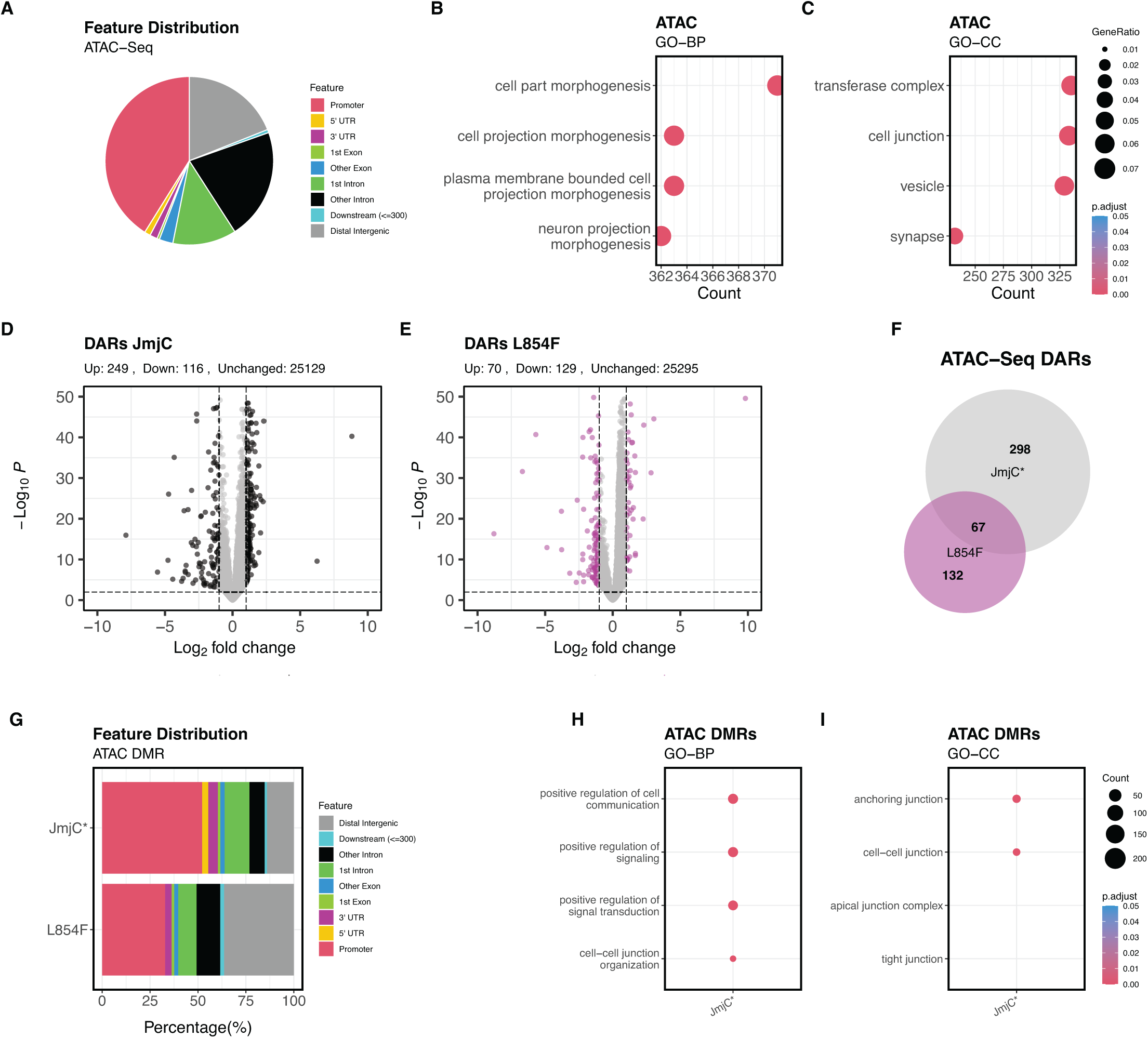
A) Pie chart showing the distribution of ATAC-Seq peaks identified in *Kdm5^WT^*. B) GO-BP of ATAC-Seq promoter-proximal peaks in *Kdm5^WT^*. C) GO-CC of ATAC-Seq promoter-proximal peaks in *Kdm5^WT^*. D) Volcano plot showing differentially accessible regions (DARs) in *Kdm5^JmjC*^* brains. Black dots indicate statistically significant changes. E) Volcano plot showing differentially accessible regions (DARs) in *Kdm5^L854F^* brains. purple dots indicate statistically significant changes. F) Venn diagram showing the overlap between DARs observed in *Kdm5^JmC*^* and *Kdm5^L854F^* brains. G) Bar plot of DARs peak annotations *Kdm5^JmC*^* and *Kdm5^L854F^* brains. H) GO-BP of ATAC-Seq DMRs in *Kdm5^JmjC*^*. I) GO-CC of ATAC-Seq DMRs in *Kdm5^JmjC*^*.

**Supplemental Figure 5.**
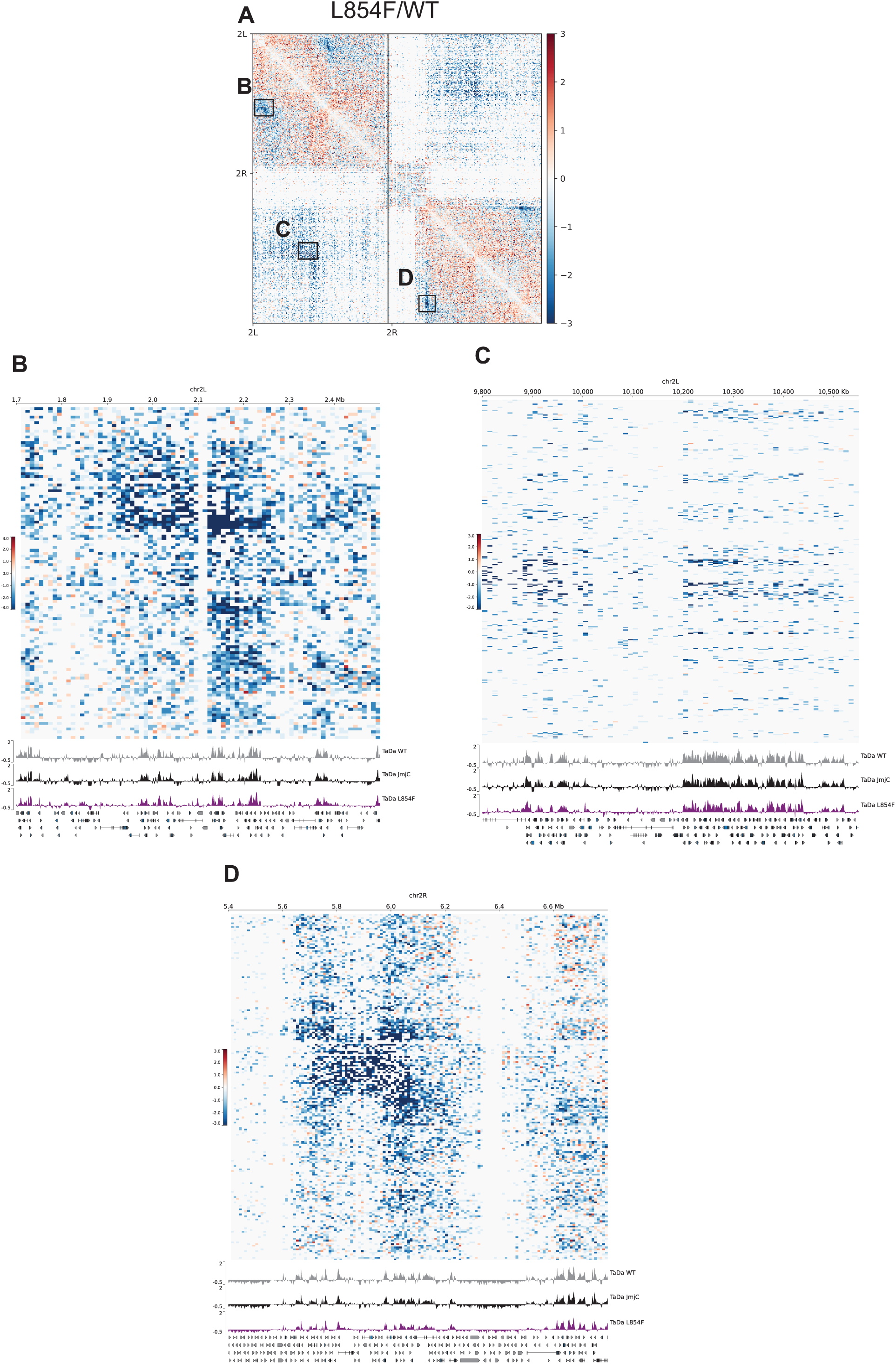
**A)** *Kdm5^L854F^*/*Kdm5^WT^* Hi-C contact map (10 kb) of chromosome 2. **B)** Zoomed in view of region indicated in panel A showing decreased interaction at loop anchor and associated binding by KDM5^WT^, KDM5^JmjC*^ and KDM5^L854F^. **C)** Zoomed in view of region indicated in panel A showing decreased interaction at loop anchor and associated binding by KDM5^WT^, KDM5^JmjC*^ and KDM5^L854F^. **D)** Zoomed in view of region indicated in panel A showing decreased interaction at loop anchor and associated binding by KDM5^WT^, KDM5^JmjC*^ and KDM5^L854F^.

